# PACT prevents aberrant activation of PKR by endogenous dsRNA without sequestration

**DOI:** 10.1101/2024.10.23.619951

**Authors:** Sadeem Ahmad, Tao Zou, Linlin Zhao, Xi Wang, Jihee Hwang, Anton Davydenko, Ilana Buchumenski, Patrick Zhuang, Alyssa R. Fishbein, Diego Capcha-Rodriguez, Aaron Orgel, Erez Y. Levanon, Sua Myong, James Chou, Matthew Meyerson, Sun Hur

## Abstract

The innate immune sensor PKR for double-stranded RNA (dsRNA) is critical for antiviral defense, but its aberrant activation by cellular dsRNA is linked to various diseases. The dsRNA-binding protein PACT plays a critical yet controversial role in the PKR pathway. We demonstrate that PACT is a direct and specific suppressor of PKR against endogenous dsRNA ligands like inverted-repeat Alu RNAs, which robustly activate PKR in the absence of PACT. PACT-mediated inhibition does not involve competition for dsRNA binding. Instead, PACT impairs PKR’s ability to scan along dsRNA––a process necessary for PKR molecules to encounter and autophoshorylate each other for activation. By scanning along dsRNA and directly interacting with PKR, PACT restricts PKR’s movement on dsRNA, reducing the likelihood of PKR molecular collisions and subsequent autophosphorylation, effectively inhibiting PKR without sequestering dsRNA. Consequently, PKR inhibition is more robust with longer and less abundant dsRNA, and minimal with abundant or short dsRNA. Thus, PACT functions to adjust the PKR activation threshold for long endogenous dsRNA without altering its inherent activity, revealing new mechanisms for establishing self-tolerance.

## INTRODUCTION

Effective antiviral defense relies on several innate immune receptors that detect viral dsRNAs and trigger antiviral immune responses^1,2^. These receptors include a protein kinase R (PKR) that inhibits global protein synthesis upon recognizing dsRNA, RIG-I-like receptors (RLRs) that activate the type I interferon signaling pathway, and oligoadenylate synthases (OASes) responsible for activating RNase L, leading to the widespread degradation of both cellular and viral RNAs^1,2^. While these dsRNA receptors work in synergy to mount effective antiviral immune responses, recent studies have brought to light the unique role of PKR as a key determinant of cellular fate in response to dsRNA, in particular endogenous dsRNA accumulating under dysregulated conditions^3–10^. Studies have also shown that PKR is a major contributor to the pathogenesis of a broad range of neurodevelopmental and auto-inflammatory diseases, including Parkinson’s disease^11–14^, systemic lupus erythematosus (SLE)^15^ and proteasome-associated autoinflammatory syndromes (PRAAS)^16^, although the underlying molecular mechanisms for the aberrant PKR activation under these disease conditions and its tissue specificity are only beginning to be understood^17,18^.

PKR represents one of the four kinases that initiate integrated stress response (ISRs), a conserved cellular response to common stressors, including oxidative stress, nutrient starvation and viral infection^19^. In the resting state, PKR is in a latent, unphosphorylated and monomeric state. Upon dsRNA binding, however, PKR undergoes dimerization and autophosphorylation, both of which cooperate to activate its kinase activity^20,21^. In this process, dsRNA serves as a scaffold to bring together PKR molecules, facilitating their dimerization and inter-molecular phosphorylation^22,23^. This notion is supported by observations that PKR activation requires a minimum of ∼30 bp-long dsRNA, sufficient to accommodate two PKR molecules, and that PKR is inhibited by an excess amount of dsRNA, which disperses PKR among many dsRNA molecules, preventing autophosphorylation between PKR molecules^23,24^. Once activated, PKR phosphorylates the translational initiation factor eIF2α, which then results in the inhibition of translational initiation for the majority of capped mRNAs except for a small group of stress-related genes that help the cell recover from stress conditions^25,26^. While transient activation of ISR is beneficial for the cell, its prolonged activation can lead to cellular toxicity through various mechanisms^19,27^.

The importance of PKR in cell fate decision is underscored by the multi-layered regulatory mechanisms for PKR. One extensively studied negative regulator is ADAR1, an RNA editing enzyme that converts adenosines (A) to inosines (I) within duplex RNA. A-to-I modification typically weakens Watson-Crick base-pairing, disrupting dsRNA structure^28^. This disruption plays a crucial role in preventing inappropriate activation of multiple dsRNA sensors in the cell, including PKR and averting unwanted innate immune system activation^5–10^. More recent studies suggested that ADAR1 can also suppress PKR in a manner independent of the catalytic activity^6^, implicating the importance of its RNA binding activity, in addition to editing functions.

PACT is another protein suggested to modulate PKR and other dsRNA sensing pathways, but its functions and mechanisms have been controversial. Widely considered a <u>P</u>KR <u>act</u>ivator^29–32^ ––thus named PACT––more recent genetic studies contradicted this prevailing notion. Knocking out the PACT homolog (RAX) in mice causes reproductive and developmental defects, which were rescued by PKR double knock-out (DKO)^33^. Impaired expression of PACT in 293T or Hela cells^34,35^ also showed increased activity of PKR. Although the genetic evidence suggested the PKR-suppressive functions of PACT^33–35^, the underlying mechanisms––including whether such effects are direct or indirect––have been unclear, largely because these inhibitory functions have not been biochemically reconstituted. In fact, earlier biochemical studies suggested that PACT may activate PKR independent of dsRNA^29,30,36^, raising questions about the exact functions of PACT and how these findings can be reconciled altogether.

In contrast to PACT’s seemingly complex and controversial biological functions, its domain architecture is remarkably simple, featuring just three tandem repeats of dsRNA-binding domains (DRBDs). DRBDs are common in dsRNA-binding proteins across all life kingdoms and viruses, with a conserved fold and mode of dsRNA interaction^37,38^. DRBDs commonly recognize dsRNA backbone structure with little dependence on dsRNA sequence^38^. However, certain DRBDs have evolved to function as protein-protein interaction domains at the expense of dsRNA-binding activity^39–42^. For PACT, the first two DRBDs exhibit typical sequence-independent dsRNA-binding activity. The third DRBD, however, shows little to no dsRNA binding activity and instead forms a homodimer^43,44^, showcasing a divergent evolution of DRBD as a protein-interaction domain.

We here examine how PACT modifies the PKR activity using a combination of biochemistry, single-molecule analyses, cellular functional studies, and structure-guided mutagenesis. Our data unambiguously indicate that PACT is a direct and specific negative regulator of PKR, restricting PKR’s movement on dsRNA and limiting its ability to undergo autophosphorylation. This reveals a unique mode of regulating foreign nucleic acid sensors in innate immunity.

## RESULTS

### PACT restricts aberrant activation of PKR, thereby maintaining cellular homeostasis

Previous studies showed that aberrant activation of PKR leads to the loss of cell fitness in many cancer cell lines. To investigate the role of PACT in PKR functions, we examined the impact of PACT depletion using 10 different cancer cell lines. In 5 out of 10 cells tested (HCC1806 - breast, MDA-MB-157 - breast, NCI-H727 - lung, NCI-H2286 - lung, SW-1271 - lung), knocking out PACT led to a loss of cell fitness, as measured by ATP bioluminescence assay (Figure 1A). We further chose three of these cell lines—HCC1806, NCI-H727, and NCI-H2286—and confirmed their sensitivity to PACT depletion using crystal violet staining, an orthogonal cell viability assay (Figure 1B). This loss of cell fitness could be prevented by overexpressing wild-type PACT in PACT-deficient HCC1806 cells (Figure 1C), further supporting the role that PACT as an essential gene. Note that for PACT overexpression, a version of wild-type PACT that is resistant to targeting by PACT sgRNA #1 was used.

**Figure 1.**
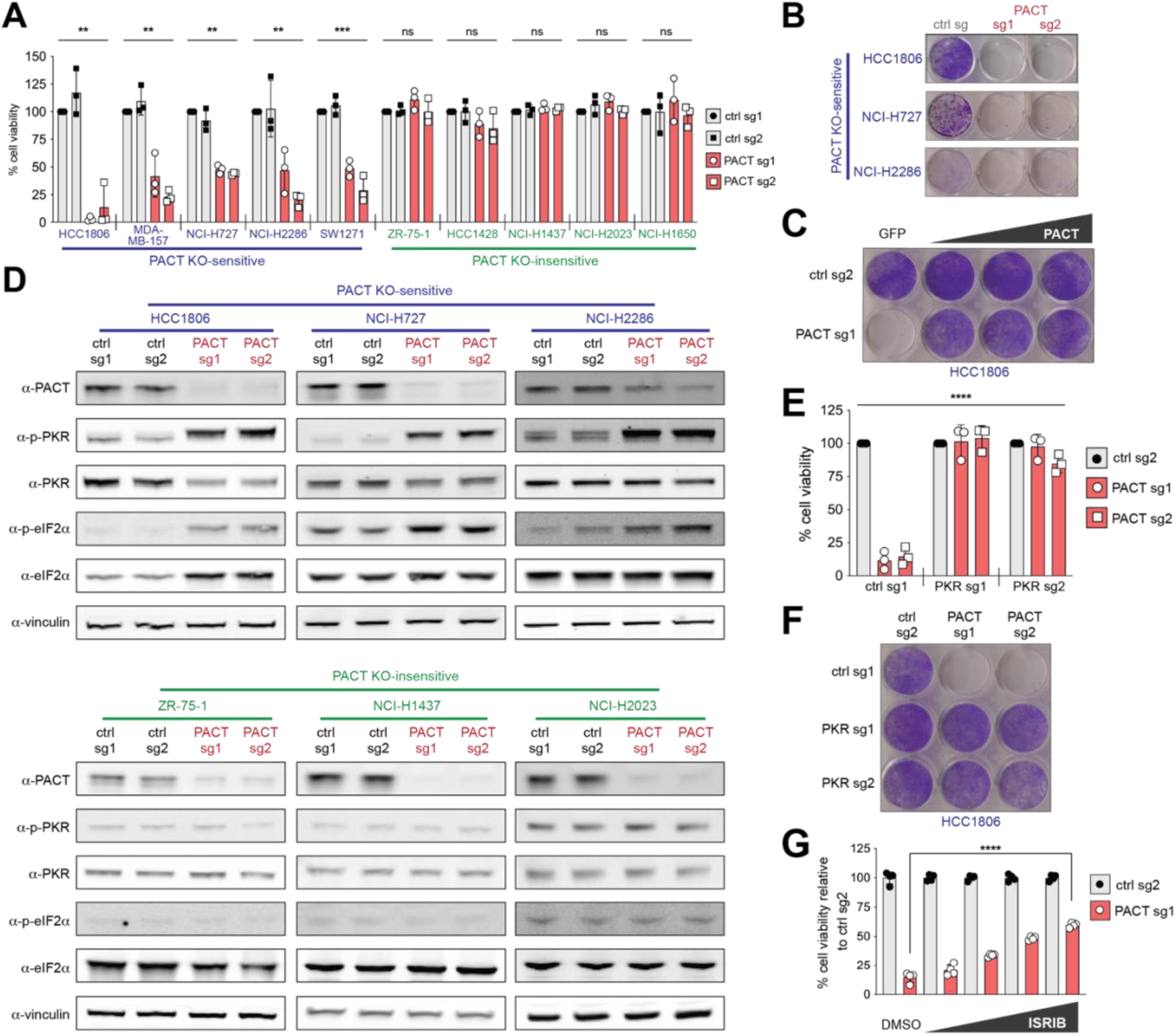
PACT restricts aberrant activation of PKR, thereby maintaining cellular homeostasis. (A) ATP bioluminescence assay showing cell viability in PACT KO-sensitive (HCC1806, MDA-MB-157, NCI-H727, NCI-H2286, SW1271) and -insensitive (ZR-75-1, HCC1428, NCI-H1437, NCI-H2023, NCI-H1650) cell lines after control gene-KO (ctrl) or PACT-KO using 2 different guide RNAs. The control (ctrl) genes are AAVS1 (sg1) and Chr2.2 (sg2). Values represent means of 3 biological repeats. *P*-values were based on one-way ANOVA test. (B) Crystal violet staining assay showing cell viability in 3 PACT KO-sensitive cells (HCC1806, NCI-H727 and NCI-H2286) after control gene-KO vs. PACT-KO. (C) Crystal violet staining assay with HCC1806 cells after control gene-KO and PACT-KO complemented with increasing amounts of wild-type PACT. PACT used here was engineered to be resistant to PACT sgRNA 1 targeting. (D) Western blot analysis showing levels of PKR and eIF2α phosphorylation in PACT KO-sensitive (top panel) and -insensitive (bottom panel) cells after control gene-KO or PACT-KO. Vinculin was used as a loading control. (E) ATP bioluminescence assay in HCC1806 cells in PKR-sufficient and PKR-KO cells in control gene-KO and PACT-KO backgrounds. Values represent means of 3 biological repeats. *P*-values were based on two-way ANOVA test. (F) A representative crystal violet staining for samples in (E). (G) ATP bioluminescence assay in HCC1806 cells in control gene-KO and PACT-KO treated with indicated concentrations of ISRIB. Values represent means of 3-4 technical repeats. *P*-values were based on two-tailed student’s t test. **** *p*= <0.0001, *** *p* = <0.001, ** *p* = <0.01, * *p* = <0.05, not significant (ns) is for *p* > 0.05

To examine whether PKR is aberrantly activated by PACT depletion, we measured the levels of phospho-PKR (p-PKR) and phospho-eIF2α (p-eIF2α) by western blot (WB). The results showed that PACT dependency segregated with PKR activation. That is, PACT depletion increased PKR activity in PACT knockout (KO)-sensitive cells, but not in PACT KO-insensitive cells (Figure 1D). Similarly, PACT depletion led to the induction of *CHOP*, a known consequence of eIF2α phosphorylation^45^, in PACT KO-sensitive cells, but not in PACT KO-insensitive cells (Figure S1A). Consistent with the notion that aberrant activation of PKR caused increased eIF2α phosphorylation and loss of cell fitness, double knockout (DKO) of PKR returned p-eIF2α to the basal level and rescued cell viability (Figures 1E, 1F, Figures S1B-D). Interestingly, ISRIB, a small molecule inhibitor of ISR^46^, showed varying degrees of partial rescue in 2 different PACT-dependent cell lines, suggesting that PKR regulates cell fitness through both ISR-dependent and -independent mechanisms (Figures 1G, S1E-F).

We then examined how other dsRNA-dependent innate immune pathways are affected by PACT depletion and whether PACT’s inhibitory function is specific to PKR. To test OAS activity, we measured the integrity of rRNAs, which are well-known targets of RNase L downstream of OASes. Bioanalyzer data showed that rRNAs are intact in all cells, suggesting minimal activation of OASes in the absence of PACT, regardless of a cell line’s sensitivity to PACT depletion (Figure S1G). Furthermore, we did not observe RLR activation, as measured by *IFNβ* and *IL-6* induction, in most cells, except for the HCC1806 cell line (Figure S1A). Consistent with these observations, knocking out the RNase L or the RLR adaptor MAVS had little impact on either cell viability or PKR activity (Figures S2A-D). Altogether, these results suggest that PACT depletion leads to aberrant activation of PKR and consequent loss of cell fitness, without necessarily activating other dsRNA-dependent innate immune signaling pathways.

### PACT shares dsRNA ligands with PKR, but does not sequester dsRNA away from PKR

We next examined how PACT inhibits PKR. Since PACT is a dsRNA-binding protein (*K*_d_ of 357.5 nM), we asked whether PACT’s dsRNA-binding activity is important. We generated the dsRNA binding-deficient variant that harbors four Lys-to-Glu mutations (4KE) in the dsRNA interface of DRBD1 (K84, K85) and DRBD2 (K177, K178) (Figure 2A). Complementation with WT PACT in PACT-depleted HCC1806 cells restored cell viability, but 4KE PACT did not (Figure 2B), indicating that PACT requires dsRNA binding to regulate PKR.

**Figure 2.**
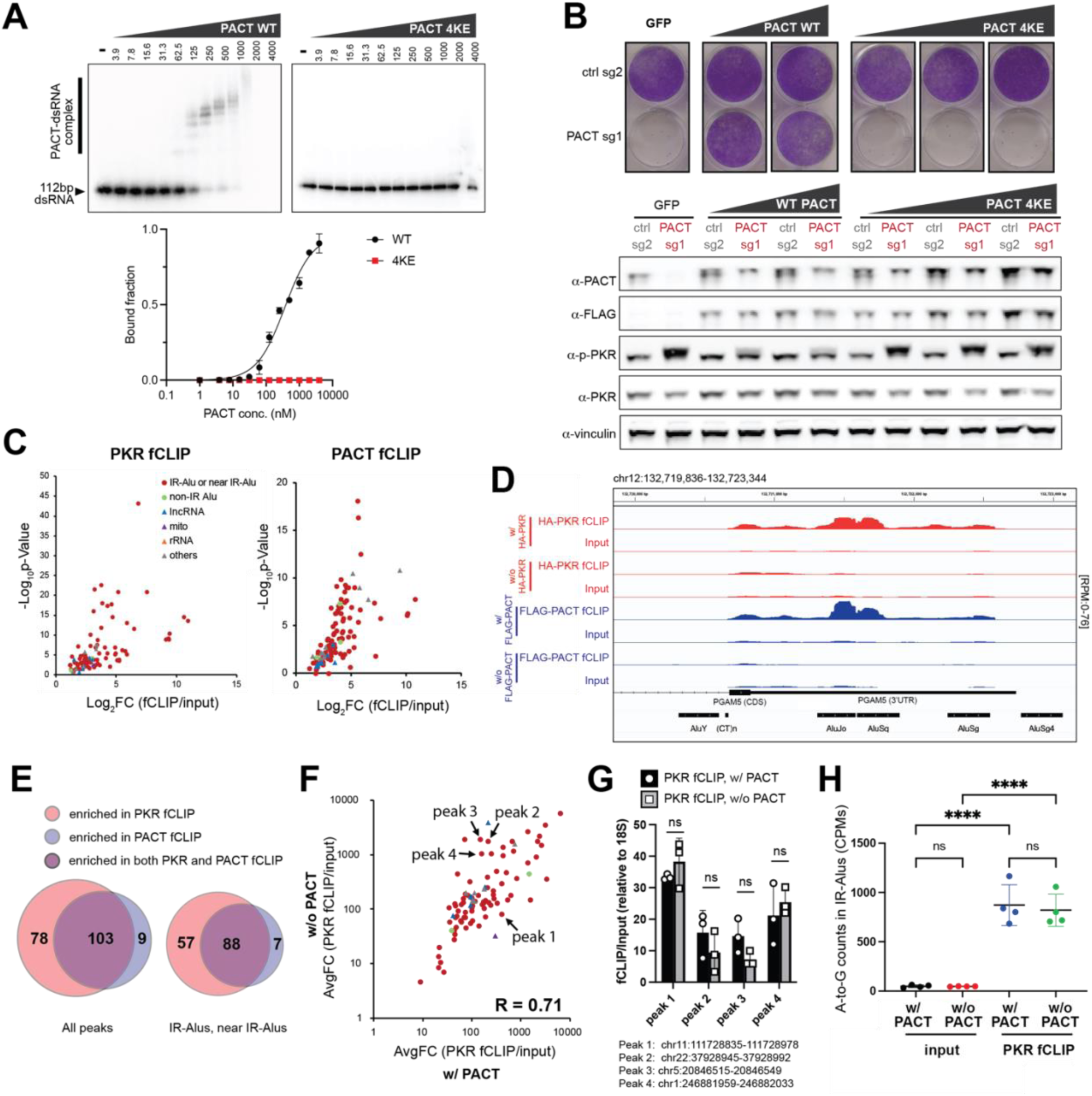
PACT and PKR share endogenous dsRNA ligands, but PACT inhibits PKR without blocking PKR’s dsRNA binding. (A) Native gel-shift assay with 112 bp dsRNA binding to increasing concentrations (0, 3.9, 7.8, 15.6, 31.3, 62.5, 125, 250, 500, 1000, 2000, 4000 nM) of PACT wild-type or 4KE (K84E, K85E, K177E, K178E) mutant. Right panel: specific binding curve using the Hill equation assuming 8 protein binding sites (see Methods). Wild-type plot represents mean ± S.D. from 3 experiments. (B) Crystal violet staining assay showing viability of HCC1806 cells after control gene-KO and PACT-KO complemented with expression of increasing amounts of PACT wild type and 4KE mutant. Below: WB analysis of the samples above. (C) Volcano plots showing RNA peaks enriched by PKR fCLIP (left) and PACT fCLIP (right). The enriched peaks were filtered using the criteria: (Log_2_FC (HA-fCLIP/input) >=1 in cells expressing HA-PKR & Log_2_FC (HA-fCLIP/input) <=0 in cells without HA-PKR) and (Log_2_FC (FLAG-fCLIP/input) >=1 in cells expressing FLAG-PACT & Log_2_FC (FLAG-fCLIP/input) <=0 in cells without FLAG-PACT). See also Figures S3C, S3F. HCC1806 cells were used in all experiments. Values represent means of 3 and 2 biological repeats for PKR fCLIP and PACT fCLIP respectively. (D) A representative snapshot of an IR-Alu sequence enriched in PKR (red) and PACT (blue) fCLIP-seq results. (E) Venn diagram of total peaks (left) and IR-Alu or near IR-Alu peaks (right) enriched in PKR and PACT fCLIP. A peak was considered near IR-Alu if it was either flanked by a pair of IR-Alus (within 300 bp) or in the vicinity of an IR-Alus (<300 bp away). (F) Comparison of HA-PKR fCLIP intensity from HCC1806 cells before (x-axis) and after PACT KO (y-axis). The plot shows the average fold change of AUC (area under curve) for each peak identified from (C). Values were obtained by first normalizing indicated RNA levels (AUC) to the total non-rRNA reads (because rRNAs were depleted during the library preparation), followed by calculating the ratio of fCLIP to input for the normalized AUC. See Figure S3G and Methods for additional discussion of the normalization. The plot represents the mean from 2 biological repeats. The correlation coefficient (R) is 0.71. (G) fCLIP-RTqPCR of 4 peaks selected from (F) using endogenous PKR. fCLIP was performed using anti-PKR. Values were obtained by first normalizing indicated RNA levels to the internal control 18S rRNA, and then calculating the ratio of fCLIP to input for the normalized RNA levels. The results represent mean ± SD from 3 independent experiments. *P*-values were calculated by two-tailed t-test. (ns, not significant; *P*>0.05). (H) Level of A-to-I editing in PKR fCLIP and input samples from HCC1806 cells before and after PACT KO. Y-axis values represent A-to-G mismatch counts per million reads (CPM). Values represent means of 4 biological repeats. *P*-values were calculated by one-way ANOVA followed by Tukey multiple comparisons test. **** *p* <0.0001; (ns) *p* >0.12

One possible mechanism is that PACT sequesters endogenous dsRNA, thereby limiting their access to PKR. To test this possibility, we examined whether PACT and PKR share their dsRNA ligands in cells, and whether the presence of PACT affects PKR–dsRNA interaction. To measure the interactions between PKR/PACT and dsRNA in cells, we employed the formaldehyde-assisted RNA-protein crosslinking and immunoprecipitation (fCLIP), a method well-established for capturing protein–dsRNA interactions that are often difficult to capture using direct UV crosslinking methods^47^.

We performed PKR fCLIP in PACT-depleted HCC1806 cells, one of the PACT KO-sensitive cell lines. Because PACT deletion results in significant compromise in cell viability, we generated a PACT/PKR DKO model and complemented it with the HA-tagged, kinase-dead PKR mutant (K296R, Figure S3A), which does not impair cell viability. Our initial validation of PKR fCLIP demonstrated significant enrichment of an *in vitro* transcribed 112 bp dsRNA transfected into the cells, exhibiting ∼500-fold increase over the input control (Figure S3B). Anti-HA showed a cleaner background signal than anti-PKR in 112 bp dsRNA enrichment (Figure S3B) and therefore was chosen for the subsequent PKR fCLIP experiments. We next proceeded with PKR fCLIP without the exogenously introduced dsRNA to examine the endogenous dsRNA ligands. As a control, we also performed fCLIP in the absence of HA-PKR. Among 116 PKR fCLIP peaks (Log2FC >=1), we computationally removed 12 peaks, predominantly rRNAs, which were also enriched by fCLIP in the absence of HA-PKR (Log2FC >0) (Figure S3C). This filtering led to 94 PKR-specific fCLIP peaks (Figure 2C), of which 79 peaks were invert-repeat Alu RNAs (IR-Alus) and regions proximal to IR-Alu pairs (within 300 bp of or flanked by IR-Alu pair) (see Figure 2C, D). This is consistent with the previous reports showing that IR-Alus are the primary endogenous dsRNAs in primates, and serve as the major ligands for PKR and other dsRNA binding proteins^4,5,48,49^. Unlike previous report showing association between PKR and sense-antisense overlapping transcripts from mitochondria in Hela cells^3^, we observed little sign of sense-antisense overlaps in mitochondrial transcripts either in our input or PKR fCLIP samples (Figure S3D). This discrepancy may reflect cell type-dependent variability in mitochondrial dsRNA accumulation, as previously noted^17^.

To identify RNA ligands for PACT, we performed a similar PACT fCLIP using HCC1806 cells ectopically expressing FLAG-tagged PACT. Anti-FLAG fCLIP also showed robust enrichment of exogenously introduced 112 bp dsRNA (Figure S3E), validating our approach. PACT fCLIP showed RNA ligands that largely overlapped with those identified from PKR fCLIP (Figure 2C-E, S3F). Specifically, IR-Alu pairs and regions proximal to IR-Alus, identified from PKR fCLIP, represented the majority of the PACT-bound RNA species (Figure 2E, see also Figure 2D). As with PKR fCLIP, we observed little enrichment of mitochondrial RNAs (Figure S3D).

Given that PACT and PKR largely share the endogenous dsRNA ligands, we investigated whether PACT competes with PKR for the shared dsRNA substrates. To address this, we compared PKR fCLIP intensity from PACT-sufficient and -deficient cells, and found that most PKR peaks were largely unaffected by PACT (Figure 2F, see Figure S3G for normalization validation). While a small subset of PKR fCLIP peaks showed variable changes in intensity upon PACT depletion, their changes were statistically insignificant as observed by fCLIP RT-qPCR (Figure S3H). Furthermore, comparison of fCLIP RT-qPCR with endogenous PKR corroborated this finding (Figure 2G).

We also examined whether ADAR1-mediated A-to-I editing of dsRNA, known to affect PKR activity, was somehow affected by PACT. However, the A-to-I editing level of either PKR fCLIPed IR-Alus or overall IR-Alus were not affected by the presence of PACT (Figure 2H, S3I).

These results suggest that PACT regulates PKR without limiting PKR’s access to endogenous dsRNA, and this regulation is independent of ADAR1. Additionally, given that PACT knockout primarily affects PKR but not ADAR1 (Figure 2H), OASes-RNase L (Figure S1G), and RLRs (in two out of three cell lines tested for RLRs) (Figure S1A), we conclude that PACT can inhibit PKR without globally sequestering cellular dsRNAs.

### PKR scans along dsRNA, resulting in dsRNA length-dependent activity

To elucidate the molecular mechanism by which PACT inhibits PKR, we employed an *in vitro* PKR activity assay. We measured PKR autophosphorylation using radiography with [γ^32^P]-ATP–– a well-established readout for PKR activity. After the kinase reaction with [γ^32^P]-ATP, PKR was analyzed on SDS-PAGE for the quantitative measurement of [^32^P]-incorporated PKR. Note that we used [^35^S]-labeled IRF3 as a loading control, allowing us to compare intensities across different gels (Figure 3A).

**Figure 3.**
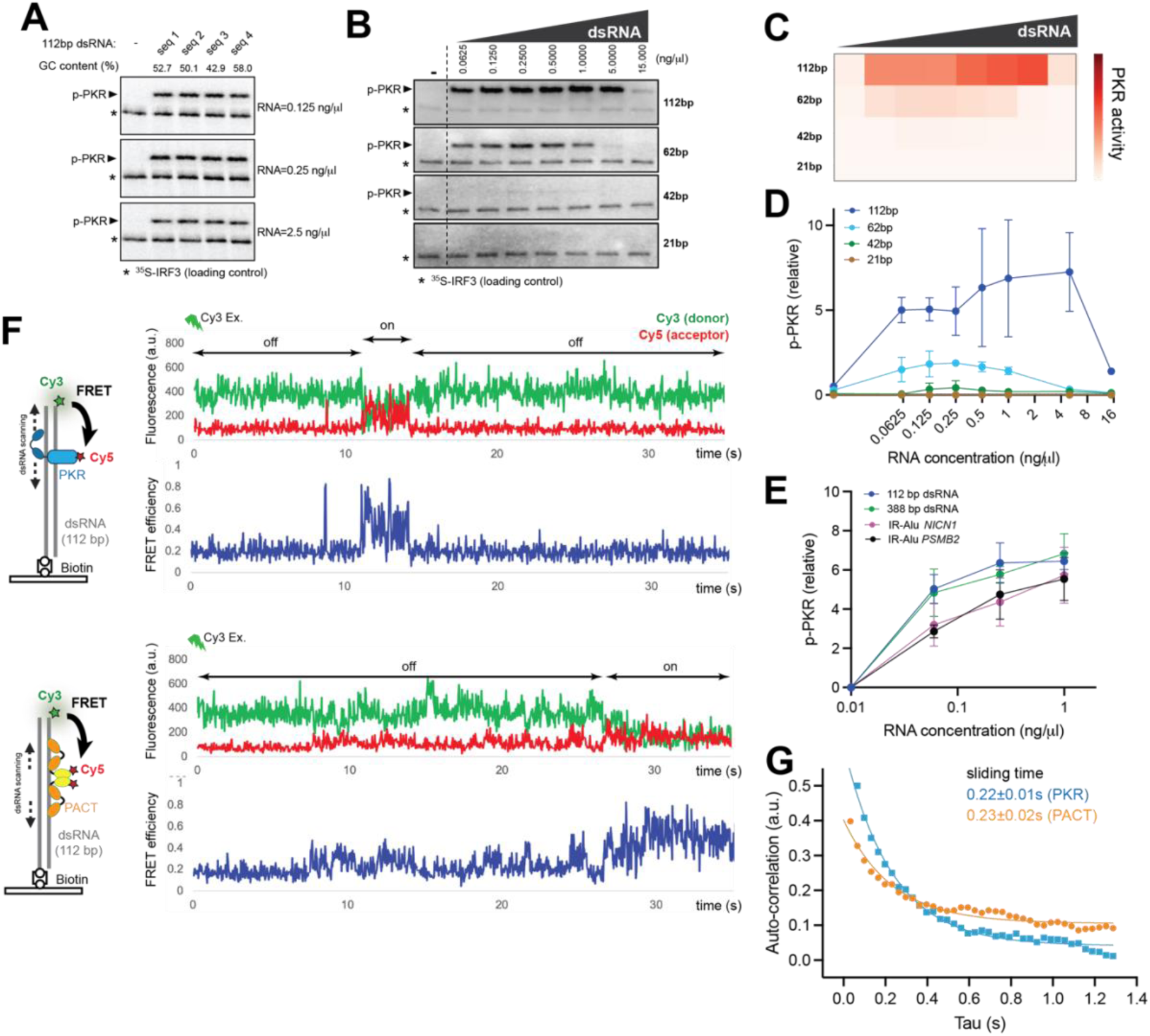
PKR scans along dsRNA, resulting in dsRNA length-dependent activity. (A) *In vitro* PKR kinase assay monitoring PKR autophosphorylation using [^32^P]-ATP in the presence of increasing concentrations (0.125, 0.25, 2.5 ng/μl) of 112 bp dsRNA with 4 different sequences (see supplementary Table 3) and GC contents. [^35^S]-IRF3 was used as a loading control (*). Unless mentioned otherwise, wild-type full-length PKR was used. (B) *In vitro* PKR kinase assay with increasing concentrations (0.0625, 0.125, 0.25, 0.5, 1, 5, 15 ng/μl) of dsRNA of lengths 112, 62, 42, 21 bp. (C-D) Heatmap and x-y graph representations of the PKR activity quantified from the gels in (A). Values represent means of 3 biological repeats. (E) PKR activity with 112 bp dsRNA, perfect duplex Alu dsRNA, IR-Alus from 3’-UTRs of *NICN1* and *PSMB2* mRNAs (see Figure S4E for secondary structures) using the RNA concentrations of 0.0625, 0.25, 1 ng/μl. Values represent means of 3 biological repeats. (F) Schematic of single molecule FRET experiments to monitor scanning motions of PKR and PACT on dsRNA. Cy5-labeled PKR or PACT (acceptor, 50 nM) was added to a chamber containing Cy3-labeled dsRNA (donor) immobilized on the surface. Cy3 and Cy5 fluorescence were monitored upon Cy3 excitation using TIRF microscopy. Time course traces of Cy3/Cy5 intensities and FRET for PKR (top) and PACT (bottom). PKR 296R was used to ensure analysis of PKR movement before its activation. (G) Autocorrelation analysis of FRET signals for PKR (blue) and PACT (orange). Sliding times were calculated by single exponential curve fitting (see Methods). Over 200 diffusion events were analyzed for each protein.

We first assessed PKR activity in the absence of PACT against various dsRNA sequences, lengths and concentrations. Consistent with previous studies^50^, PKR showed little dependence on dsRNA sequence or GC content (Figure 3A). In contrast, we observed a significant dependence on dsRNA length. Although PKR activity is known to require dsRNA lengths greater than 30 bp— the minimal length required for simultaneous occupancy by two PKR molecules^23^—we observed only weak activity with 42 bp dsRNA. This was despite confirming the monomeric PKR footprint of 15 bp (Figures S4A and S4B). In comparison, 62 bp dsRNA showed robust activation, and 112 bp dsRNA produced an even stronger response (Figures 3B–D). This was true whether PKR activity was compared at equivalent RNA mass concentrations (Figure 3D) or at molar concentrations (Figure S4C). Most importantly, increasing the dose of shorter dsRNAs, such as 62 bp, could not elevate PKR activity to levels comparable to those observed with 112 bp dsRNA (Figure 3D).

This length dependence was more evident when there was an excess amount of dsRNA. For instance, at 5 ng/µl dsRNA––corresponding to 620 nM PKR binding sites (i.e. 15 bp regions) regardless of dsRNA length, which is a significant excess over the 100 nM PKR used in our assay––PKR activity persisted nearly at peak levels with 112 bp dsRNA, whereas with 42 and 62 bp dsRNA, PKR activities were barely above the detection limit (Figures 3C-3D). This resulted in a flatter bell-shaped dsRNA dose curve with 112 bp dsRNA, compared to those with 42 and 62 bp dsRNA, allowing PKR to be activated by longer dsRNA in a manner less sensitive to the precise ratio between PKR and dsRNA.

We next asked how PKR is activated by its endogenous ligands IR-Alus, and how structural irregularities, such as mismatches or bulges, present within IR-Alus affect PKR activity. Comparing 112 bp dsRNA and 388 bp perfect Alu dsRNA (without structural irregularities) showed similar levels of PKR activity (Figure 3E), suggesting that PKR length sensitivity saturates around 100-400 bp. IR-Alus derived from the 3’UTRs of *NICN1* and *PSMB2* genes, which harbor ∼20% mismatches and bulges (Figure S4D), were less efficient at activating PKR than perfect dsRNA of the same length (Figure 3E), suggesting that PKR activity is reduced by mismatches and bulges. However, IR-Alus still exhibited over 60% of the PKR activation seen with 388 bp perfect dsRNA across all RNA concentrations tested, supporting the potential of IR-Alus to potently stimulate PKR.

One possible explanation for the observed dsRNA length-dependent activity of PKR could be multimerization along dsRNA length, similar to what has been observed with another dsRNA sensor, MDA5^51,52^, which also displays dsRNA length-dependent activity. By negative stain EM, MDA5 showed filament along the length of dsRNA, but PKR did not show similarly stable multimerization (Figure S4E). An alternative explanation for dsRNA length-dependent PKR activation is the one-dimensional diffusion of PKR along dsRNA, a behavior seen with many nucleic acid-binding proteins, especially those with tandem repeats of DRBDs, such as PACT and TRBP^53,54^. Such a scanning activity could facilitate transient encounters between PKR molecules on the same dsRNA molecule, thereby promoting *in-trans* autophosphorylation. It could also explain why PKR tolerates an excess amount of longer dsRNA than shorter dsRNA. This is because PKR molecules dispersed along longer dsRNA can interact through their diffusion along the dsRNA, whereas such interactions occur less effectively for PKR molecules dispersed on separate short dsRNAs.

To investigate whether PKR can indeed scan along dsRNA, we conducted single-molecule fluorescence resonance energy transfer (FRET) measurements between Cy3-labeled 112 bp dsRNA (donor) and Cy5-labeled PKR (acceptor) (Figure 3F, left). The dsRNA was immobilized on a surface via a biotin label, and PKR was introduced. We utilized a catalytically inactive K296R PKR to ensure analysis of PKR movement before its activation. At a concentration of 50 nM, PKR bound to dsRNA infrequently, typically exhibiting one or two binding events over approximately 30 seconds (Figure 3F). However, during the bound state, which lasted around 12.5 seconds on average (“on” duration, Figure 3F), we observed rapid fluctuations in FRET, similar to patterns previously observed with TRBP and PACT^53,54^. The clustering of these rapid fluctuations within specific time blocks, along with their infrequent occurrence, suggests that the fluctuations are unlikely due to rapid association and dissociation of PKR from dsRNA. Instead, they are consistent with rapid scanning motion along the dsRNA. Similar behavior was also observed with Cy5-labeled PACT, as previously reported^53,54^ (Figure 3F). To compare the kinetics of this scanning motion, we performed autocorrelation analysis, a common method for examining rapid diffusion behavior. Both PKR and PACT exhibited comparable scanning activities, with the sliding time corresponding to 0.22 and 0.23 seconds, respectively (Figure 3G). These observations indicate that PKR can indeed scan along dsRNA, providing a rationale for the observed dsRNA length- and concentration-dependent activity of PKR.

### PACT inhibits PKR by restricting its motion on dsRNA, leading to length-dependent inhibition

We next examined how PACT affects PKR activity, analyzing its effect across various dsRNA lengths and concentrations. For simplicity, we visualized the PKR activity as a series of heatmaps with increasing concentrations of PACT (Figure 4A, 4B, see Figure S5A for graphs). The addition of PACT raised the dsRNA threshold for PKR for all tested dsRNA lengths, confirming PACT’s role as an inhibitor rather than an activator. Similar inhibitory activities were observed with four different dsRNAs with varying sequences and GC-contents as well as IR-Alus (Figure S5B, S5C). We note that a robust and reproducible inhibitory effect of PACT was observed only after implementing an elaborate purification process to eliminate contaminating RNA from the bacterial expression system. We also confirmed that there is no detectable nuclease contamination in purified PACT as measured by the integrity of both dsRNA/ssRNA co-incubated with PACT for 2 hr at 37°C (Figure S5D).

**Figure 4.**
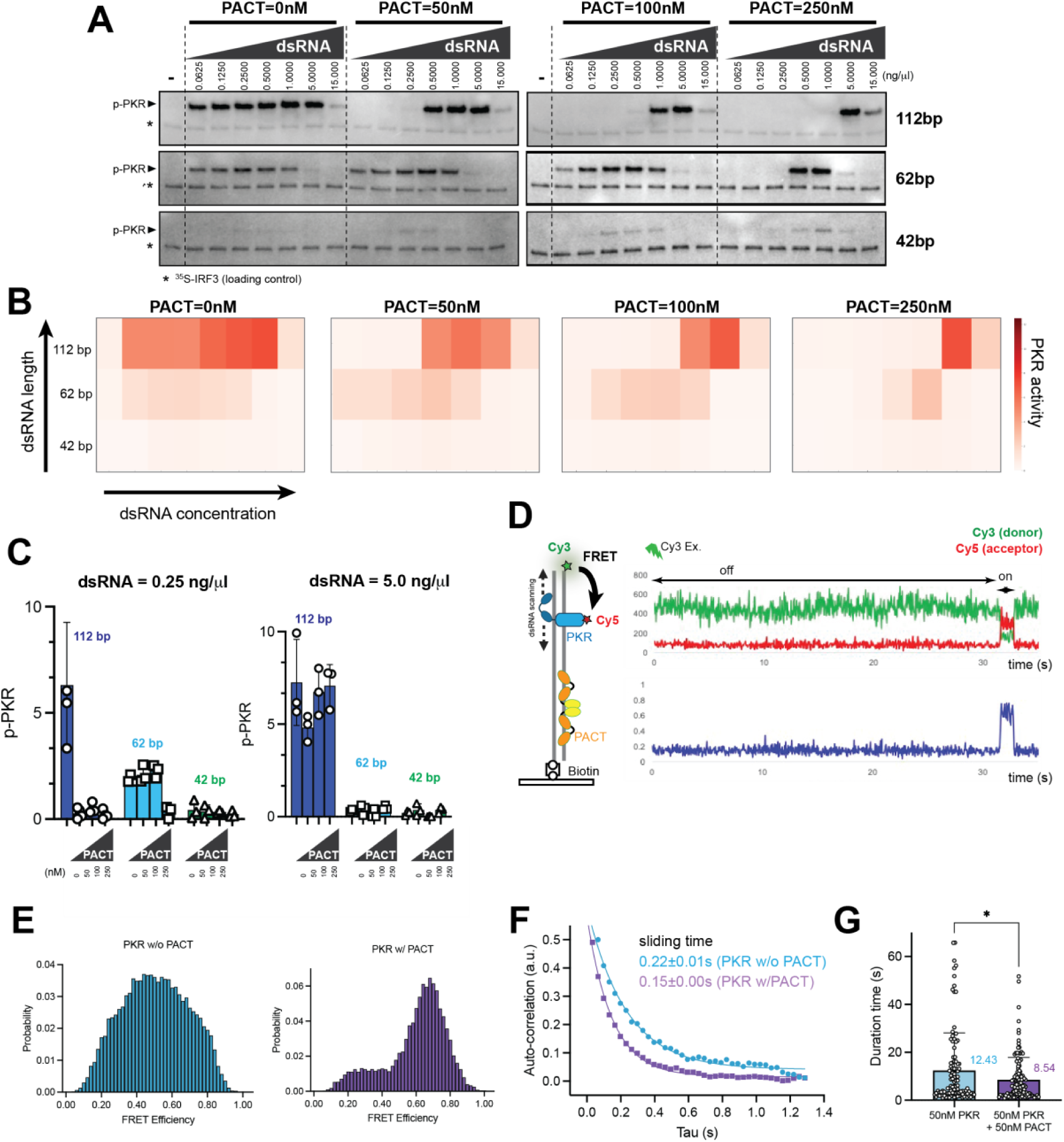
PACT inhibits PKR by restricting its motion on dsRNA, leading to length-dependent inhibition. (A) *In vitro* PKR kinase assay as in Figure 3B, but in the presence of increasing concentrations of PACT (0, 50, 100, 250 nM). [^35^S]-IRF3 was used as a loading control (*). (B) Heatmap showing relative PKR activity quantified from the gels in (A). Values represent means of 3 biological repeats. (C) Relative PKR activity with 0.25 ng/μl (left panel) and 5 ng/μl (right panel) of 112, 62, 42 bp dsRNA in the presence of 0, 50, 100, 250 nM PACT. Values represent means of 3 biological repeats. (D) Schematic of single molecule FRET experiments to monitor diffusion motions of PKR on dsRNA in the presence of PACT. Cy5-labeled PKR (acceptor, 50 nM) was mixed with unlabeled PACT (50 nM) and was added to a chamber containing Cy3-labeled dsRNA (donor) immobilized on the surface. Cy3 and Cy5 fluorescence and Cy3-Cy5 FRET were monitored upon Cy3 excitation using TIRF microscopy. PKR 296R was used to ensure analysis of PKR movement before its activation. (E) FRET histograms for PKR without PACT (blue) and PKR with PACT (purple). To compare the range of PKR motions with and without PACT, we focused our analysis on traces showing a high FRET state (>0.8). Without PACT, PKR exhibits diffusion across a broad FRET range from 0.2 to 0.8, while its movement is more restricted with PACT, showing FRET values between 0.6 and 0.8. Details on histogram quantification can be found in the Methods section. (F) Autocorrelation analysis of FRET signal for PKR without PACT (blue) and with PACT (purple). Scanning times were calculated by single exponential curve fitting (see Methods). Over 200 diffusion events were analyzed for each condition. (G) Residence time of PKR on dsRNA (“on” time) in the presence and absence of PACT (50 nM). Average values (in seconds) are shown on the right. Over 200 diffusion events were analyzed for each condition.

We then explored various potential mechanisms by which PACT might inhibit PKR, testing against our data. Firstly, we considered whether PACT might constitutively bind and inhibit PKR independently of dsRNA. If this were the case, PACT’s effect on PKR would not vary with dsRNA length or concentration. Contrary to this prediction, PACT’s inhibition was strongly dependent on both dsRNA length and concentration (Figure 4B). For example, with 50 nM PACT, PKR activity with 112 bp dsRNA was nearly completely abrogated, whereas PKR activity with 42 or 62 bp dsRNA was minimally affected, requiring an increase to 250 nM PACT for notable inhibition (Figure 4C). Furthermore, PACT potently inhibited PKR activation by 112 bp dsRNA when dsRNA was present at 0.25 ng/μl, but not at 5 ng/μl (Figure 4C). This again suggests that PACT’s actions on PKR are likely mediated through its interaction with dsRNA, consistent with the essential role of PACT’s RNA binding activity in PKR inhibition (Figures 2A, 2B).

Next, we asked if PACT inhibits PKR by sequestering dsRNA away from PKR. If this were the case, its impact would not become apparent until PACT concentrations were high enough to stably occupy a significant portion of the dsRNA. However, even at 50 nM PACT––where less than 10 % of PKR-binding sites are stably occupied by PACT (Figure 2A)––PKR activity with 112 bp dsRNA (0.25 ng/µl) decreased by more than 90% (Figures 4C), suggesting that PACT’s inhibition of PKR activity does not require dsRNA sequestration, consistent with our in vivo observation in Figure 2.

We then investigated whether PACT affects PKR’s diffusion along dsRNA, thereby interfering with its self-encounters on dsRNA and subsequent autophosphorylation. Using Cy5-labeled PKR and FRET experiments as described in Figure 3F, we assessed the impact of PACT (unlabeled) on PKR scanning motions. In the presence of 50 nM PACT, PKR still displayed dsRNA binding (FRET increase) and scanning (rapid FRET fluctuations) (Figure 4D), consistent with our interpretation above that PACT (at 50 nM) should not block PKR–dsRNA binding. However, we found a reduced range of FRET fluctuations in the presence of PACT (Figure 4D), suggesting a diminished scanning range. To differentiate between motions restricted to low-, mid-, or high-FRET states within a binding event and those fluctuating across low and high states, we focused on binding events containing the high FRET state (FRET >0.8) and generated FRET histograms. In the absence of PACT, PKR exhibited diffusion across a broad FRET range (0.2 to 0.8), indicating wide movement (Figure 4E). In contrast, with PACT present, the movement was confined to FRET states between 0.6 and 0.8 (Figure 4E). Autocorrelation analysis further revealed that PACT reduced PKR’s sliding time, consistent with restricted scanning motion (Figure 4F). Moreover, the residence time of PKR on dsRNA was also shortened, albeit slightly (Figure 4G). Altogether, these results suggest that PACT restricts PKR’s movement on dsRNA, which would decrease the likelihood of PKR molecules encountering one another and undergoing autophosphorylation, thereby inhibiting PKR.

### PACT’s inhibition of PKR involves weak but direct protein-protein interactions

To understand PACT’s mechanism further, we asked whether a mere dsRNA-binding activity is sufficient to inhibit PKR. We compared PACT with other dsRNA binding proteins. They were PACT’s closest paralog TRBP and dsRNA-binding regions of RIG-I (ΔCARD) and ADAR1 (DRBD1,2,3). The results showed that TRBP inhibited PKR similar to PACT, but RIG-I and ADAR1 did not under the equivalent condition (Figure 5A). This is despite the fact that RIG-I and ADAR1 bind dsRNA with comparable affinities (Figures 5B, S6A). These observations suggest that a mere dsRNA binding activity of PACT may be insufficient for PKR inhibition, implicating a direct and specific interaction with PKR. Consistent with this prediction, we observed a direct interaction between purified PACT and GST-PKR in the absence of RNA, as measured by GST pull-down (Figure 5C). The affinity appeared low as co-purified PACT could not be detected by Coomassie or Krypton staining, requiring more sensitive western blotting. We also observed their interaction using an orthogonal approach involving microscale thermophoresis, but their affinity in the absence of dsRNA was again too low to be accurately determined (Figure S6B).

**Figure 5.**
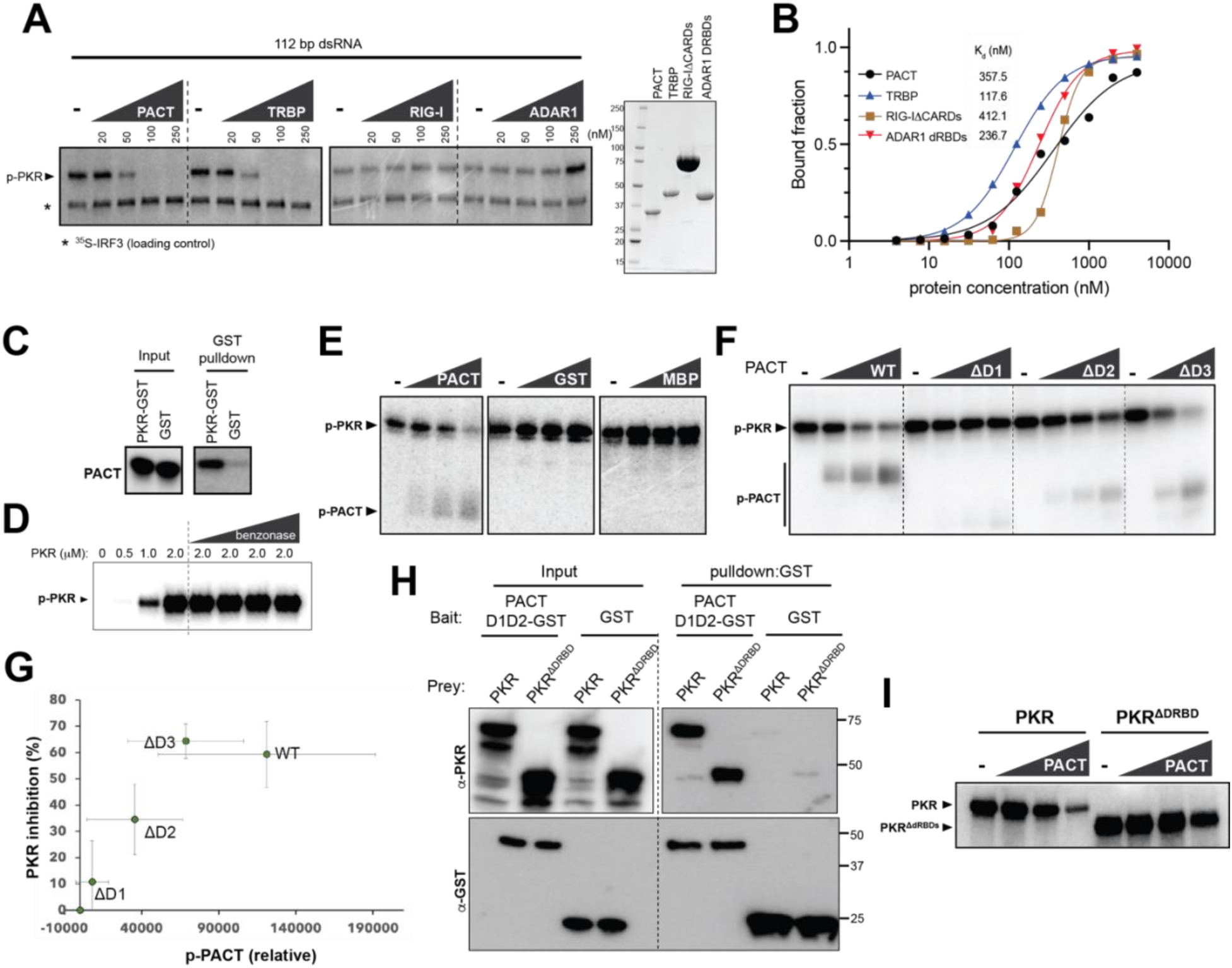
PACT’s inhibition of PKR involves weak but direct and specific protein-protein interactions. (A) *In vitro* PKR kinase assay using 112 bp dsRNA (0.25 ng/μl) in the presence of 0, 20, 50, 100, 250 nM PACT, TRBP, RIG-I ΔCARDs, ADAR1 DRBDs. Right: SDS-PAGE showing purity of the proteins. (B) dsRNA binding curve of PACT, TRBP, RIG-I ΔCARDs, ADAR1 DRBDs, derived from the native gel-shift assay in Figure S6A monitoring 112 bp dsRNA binding. For curve fitting, Hill equation was used with the assumption of 8 protein binding sites per 112 bp dsRNA (see Methods). (C) Pulldown of PACT using PKR-GST or GST. All proteins were purified and treated with benzonase to remove potential nucleic acid contaminants. Catalytically dead K296R PKR was used. (D) *In vitro* PKR kinase assay using an increasing concentration of PKR in the absence of dsRNA. Benzonase was added during the reaction (in addition to during purification) to ensure that the observed PKR activity is RNA-independent. (E) RNA-independent PKR kinase assay with 1 μM PKR in the presence of 0, 1.25, 2.5, 5 μM PACT, GST, MBP. (F) RNA-independent PKR kinase assay with 1 μM PKR in the presence of 0, 1.25, 2.5, 5 μM PACT wild type, ΔD1, ΔD2; and 0, 2.5, 5 μM ΔD3. (G) Correlation between the levels of PKR inhibition and PACT phosphorylation from (F). Values are means of 3 biological replicates. (H) Pulldown of PKR or PKRΔDRBDs using PACT D1D2-GST or GST. Catalytically dead K296R mutant was used for both PKR and PKRΔDRBDs. (I) RNA-independent PKR kinase assay with 1 μM PKR or 2 μM PKRΔDRBDs in the presence of 0, 1.25, 2.5, 5 μM PACT. Higher concentration of PKRΔDRBDs was due to its lower activity.

We hypothesized that the PACT-PKR’s direct interaction may be facilitated by their co-occupancy on dsRNA. Such RNA-facilitated protein-protein interaction is ubiquitous and can be recapitulated in the absence of RNA at high protein concentration, which bypasses the RNA requirement. Thus, we tested whether PACT could inhibit PKR activity independent of RNA at high protein concentrations. This hypothesis was testable because PKR activation can also bypass the dsRNA requirement at high PKR concentration (Figure 5D). We confirmed that the PKR activity at high protein concentration was not due to potential contaminating nucleic acids, as the addition of increasing amounts of benzonase had little impact on PKR under these conditions (Figure 5D). However, the addition of 1-5 μM PACT inhibited PKR in a dose-dependent manner (Figure 5E), suggesting that PACT can inhibit PKR independent of dsRNA at high concentrations of both PKR and PACT. Unrelated proteins, such as GST and MBP, did not show a similar PKR-inhibitory activity at equivalent concentrations (Figure 5E). Notably, PACT’s inhibition of PKR was accompanied by PACT’s phosphorylation by PKR. No such phosphorylation was observed with GST and MBP, although fewer surface-exposed Ser/Thr residues were present with these proteins (9 and 20 for GST and MBP, compared to 43 in PACT). Since phosphorylation requires direct interaction between a kinase and its substrate, this result further supports the notion that PACT forms a specific interaction with PKR.

We next examined which domains of PACT are involved in PKR suppression. In order to uncouple the requirements for dsRNA binding and PKR suppression, we continued using the assay condition with high PKR and PACT concentrations in the absence of dsRNA. Deletion of DRBD1 or DRBD2 of PACT, which are important for dsRNA binding (Figure S6C), had a significant impact on PACT’s PKR-suppressive activity (Figure 5F), whereas deletion of DRBD3 of PACT had a less significant impact on either dsRNA binding or PKR suppression (Figures S6C, 5F).

Interestingly, we observed an inverse correlation between the levels of p-PKR and p-PACT (Figure 5F). In other words, the more effectively a PACT variant inhibits PKR, the more susceptible that variant is to being phosphorylated by PKR (Figure 5G). This is the opposite of what would be expected if phosphorylation were non-specific. Furthermore, the varying levels of phosphorylation by PKR did not simply correlate with the number of surface-exposed Ser/Thr residues (43, 28, 34, and 25 for PACT, ΔDRBD1, ΔDRBD2, and ΔDRBD3, respectively), further arguing against non-specific phosphorylation. Instead, the observed correlation between PKR-mediated phosphorylation and PACT’s ability to inhibit PKR suggests a specific interaction between the two proteins that is important for PACT’s inhibitory mechanism.

Given the importance of DRBD1 and DRBD2 in PKR inhibition, and the less significant role of DRBD3, we asked whether DRBD1-2 of PACT can interact with PKR. Consistent with this notion, PKR was pulled down by PACT DRBD1-2 in the absence of RNA (Figure 5H). PACT also interacted with PKRΔDRBD (Figure 5H) and inhibited the kinase activity of PKRΔDRBD (Figure 5I), albeit not to the same extent as full-length PKR. These results together suggest that PACT can directly interact with PKR for the inhibition, and this interaction involves PACT DRBD1-2 and PKR kinase domain.

### PACT DRBD2 utilizes a surface distinct from the dsRNA interface to inhibit PKR

Our effort to further characterize the interaction between PKR and PACT using structural biology approaches proved challenging, likely due to the weak nature of binding and/or heterogeneous conformational states. Alphafold predictions were also unsuccessful. Therefore, we explored the interaction using a mutagenesis approach. To narrow down the potential interface, we surveyed other DRBDs known to form stable complexes with various protein partners and with known complex structures. These included TRBP DRBD3 in complex with Dicer^39^, TRBP DRBD3 in complex with RPAP3^41^, and Staufen DRBD5 in complex with Miranda^40^. We also determined the NMR structure of PACT DRBD3 homodimer, which revealed that two DRBD3 join their β-sheets (β1-β2-β3) through a parallel β3:β3 interaction (Figure 6A inset, Table S1).

**Figure 6.**
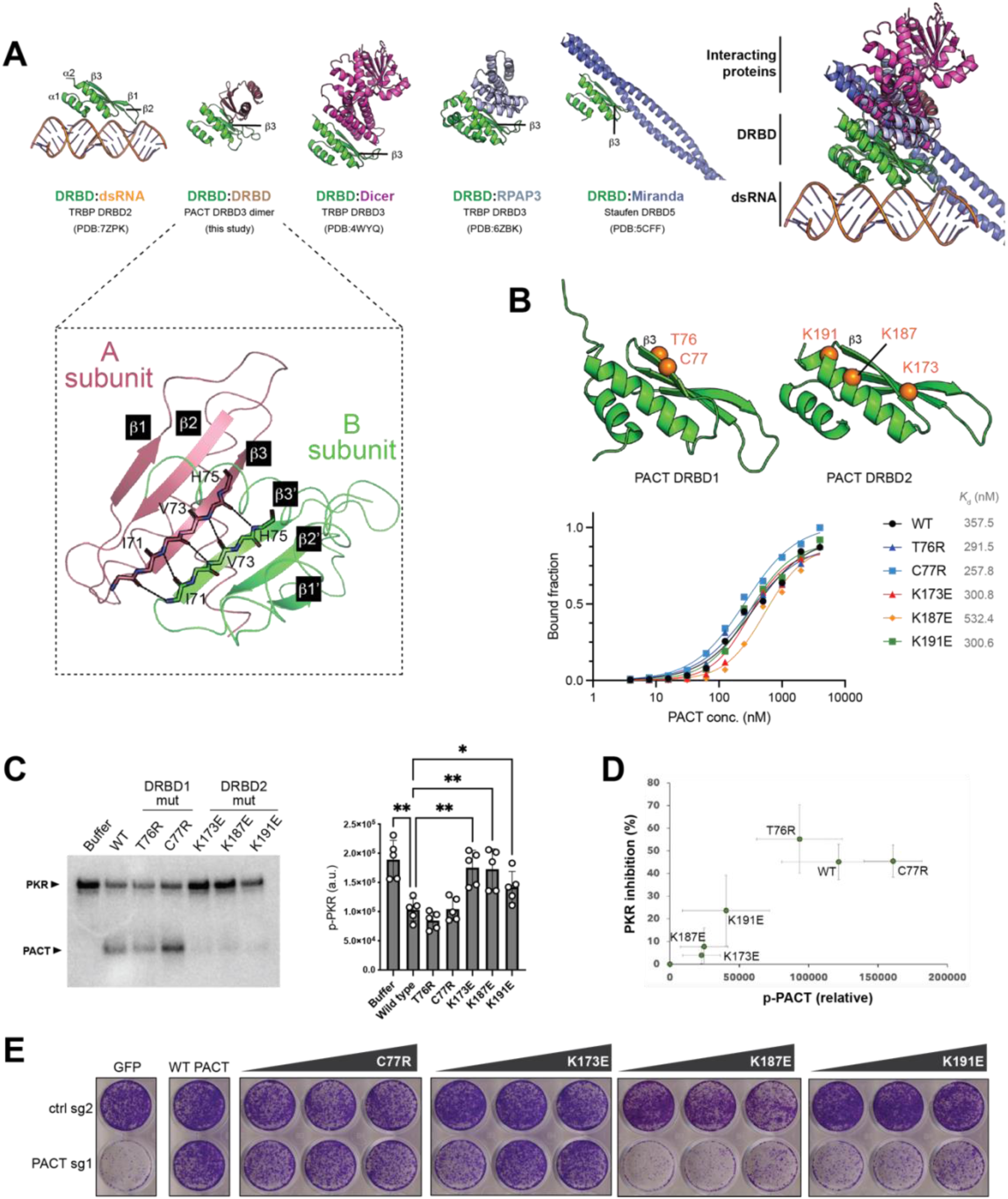
PACT DRBD2 utilizes the surface distinct from the dsRNA interface to inhibit PKR. (A) Structures of DRBDs in complex with dsRNA or protein partners, viewed from the same perspective relative to the DRBDs. Structure of PACT DRBD3 dimer is from this study. All other structures were previously published. The comparison shows the conserved protein:protein interface involving *β*3 away from the dsRNA binding surface. Inset: NMR structure of PACT DRBD3 showing its homo-dimerization by joining their β-sheets (β1-β2-β3) through a parallel β3:β3 interaction. (B) Sites of mutations on the putative protein:protein interface of PACT DRBD1 and DRBD2. Bottom: dsRNA binding curves for PACT wild type, DRBD1 mutants T76R, C77R and DRBD2 mutants K173E, K187E, K191E, derived from EMSA in Figure S6D. (C) RNA-independent PKR kinase assay with 1 μM PKR in the presence of 5 μM PACT wild type, T76R, C77R, K173E, K187E, K191E. Right: quantification from 5 independent experiments. *P*-values were calculated by one-way ANOVA followed by Tukey multiple comparisons test. ** *p* <=0.0021, * *p* <=0.0332; (ns) *p* >0.12 (D) Correlation between the levels of PKR inhibition and PACT phosphorylation from (C). Values are means of 5 biological repeats. (E) Crystal violet staining assay showing cell viability in NCI-H727 cells after control gene-KO (ctrl) and PACT-KO complemented with expression of PACT wild type or increasing amounts of C77R, K173E, K187E, K191E.

Comparison of these structures showed a striking commonality among all of these interactions– –they all utilize the same face of the DRBD opposite from the surface that is equivalent to the dsRNA interface in the canonical DRBD (Figure 6A). In particular, all utilize β3 as a key component in the protein interactions (Figure 6A). Based on these observations, we asked whether PACT also inhibits PKR through β3 of DRBD1 and DRBD2.

We generated PACT variants harboring mutations in and around β3 of DRBD1 (T76R, C77R) and DRBD2 (K173E, K187E, K191E) (Figure 6B). All mutants showed comparable affinities for dsRNA as WT PACT (Figures 6B, S6D), suggesting that they are folded correctly. This is also consistent with the structure that these residues are far from the dsRNA interface. All three DRBD2 mutations, however, impaired PKR-inhibitory activity of PACT (Figure 6C). The DRBD1 mutations did not affect the PKR-inhibitory function of PACT, suggesting different surface utilization (see the discussion in Figure S6E, S6F). We again observed the correlation between PKR’s ability to phosphorylate PACT and PACT’s inhibitory effect on PKR (Figure 6D), further supporting the notion that PACT exerts PKR inhibition through direct protein-protein interaction. Finally, we found that the DRBD2 mutants K187E and K191E were defective in restoring cell viability and inhibiting the PKR activity in PACT-dependent cells in our complementation assay (Figures 6E, S7A-S7B). In contrast, C77R was comparable to WT PACT in both the *in vitro* PKR activity assay and cellular assays (Figures 6C, 6E, S7A). Interestingly, K173E, which showed an impaired PKR-inhibitory effect *in vitro*, did not exhibit a significant defect in the cellular assays (Figures 6C, 6E, S7A), suggesting a potential compensatory mechanism in cells. Nevertheless, our mutagenesis data on K187E and K191E support the notion that PACT inhibits PKR through a direct protein-protein interaction, involving the PACT DRBD2 surface analogous to other DRBD-protein interfaces.

## DISCUSSION

PKR is a conserved sensor for foreign dsRNA responsible for activating the integrated stress response during viral infection and contributing to the pathogenesis of a broad range of immune diseases and neurodysregulation^11–15^. Despite its importance, the mechanisms regulating PKR to prevent aberrant activation by endogenous dsRNA are only beginning to be understood. Through mechanistic studies, we here demonstrate that PACT, whose functions have been controversial, plays an essential role in restraining PKR and preventing its inappropriate activation by endogenous dsRNA formed by Alu elements. Our biochemical analyses suggest that previous reports of PACT-dependent activation of PKR^29–32^ may have resulted from incomplete RNA removal from PACT, which we found requires extensive nuclease treatment and elaborate purification steps.

Our data suggest that PACT inhibits PKR through an unusual mechanism dependent on dsRNA length and concentration (Figure 7A). Although PACT and PKR share the endogenous dsRNA ligands, primarily IR-Alus, PACT does not block PKR binding to these dsRNAs per se. Instead, a substoichiometric amount of PACT binding to dsRNA is sufficient to inhibit the bulk of PKR activity. Single molecule analyses suggest that both PKR and PACT scan along dsRNAs––a characteristic common to proteins with tandem repeats of nucleic acid-binding domains^53–55^. PKR utilizes its scanning ability to encounter other PKR molecules for autophosphorylation, explaining its preference for long dsRNAs (Figure 7B). Conversely, PACT scans dsRNA to interact with PKR and restricts PKR’s movement on dsRNA. This decreases the likelihood of PKR molecules encountering one another and undergoing autophosphorylation, thereby inhibiting PKR without sequestering dsRNA (Figure 7B). Consequently, PACT’s inhibition of PKR is more robust when dsRNA is longer and less abundant (Figure 7A)––conditions that favor co-occupancy of PACT and PKR on the same RNA. This suggests that PACT’s effect on viral infection may also be virus- and dose-dependent, as different viruses generate dsRNAs of varying lengths and amounts. Collectively, PACT functions as a threshold setter––selectively raising the tolerance threshold for longer dsRNAs without compromising PKR’s inherent activity (Figure 7A).

**Figure 7.**
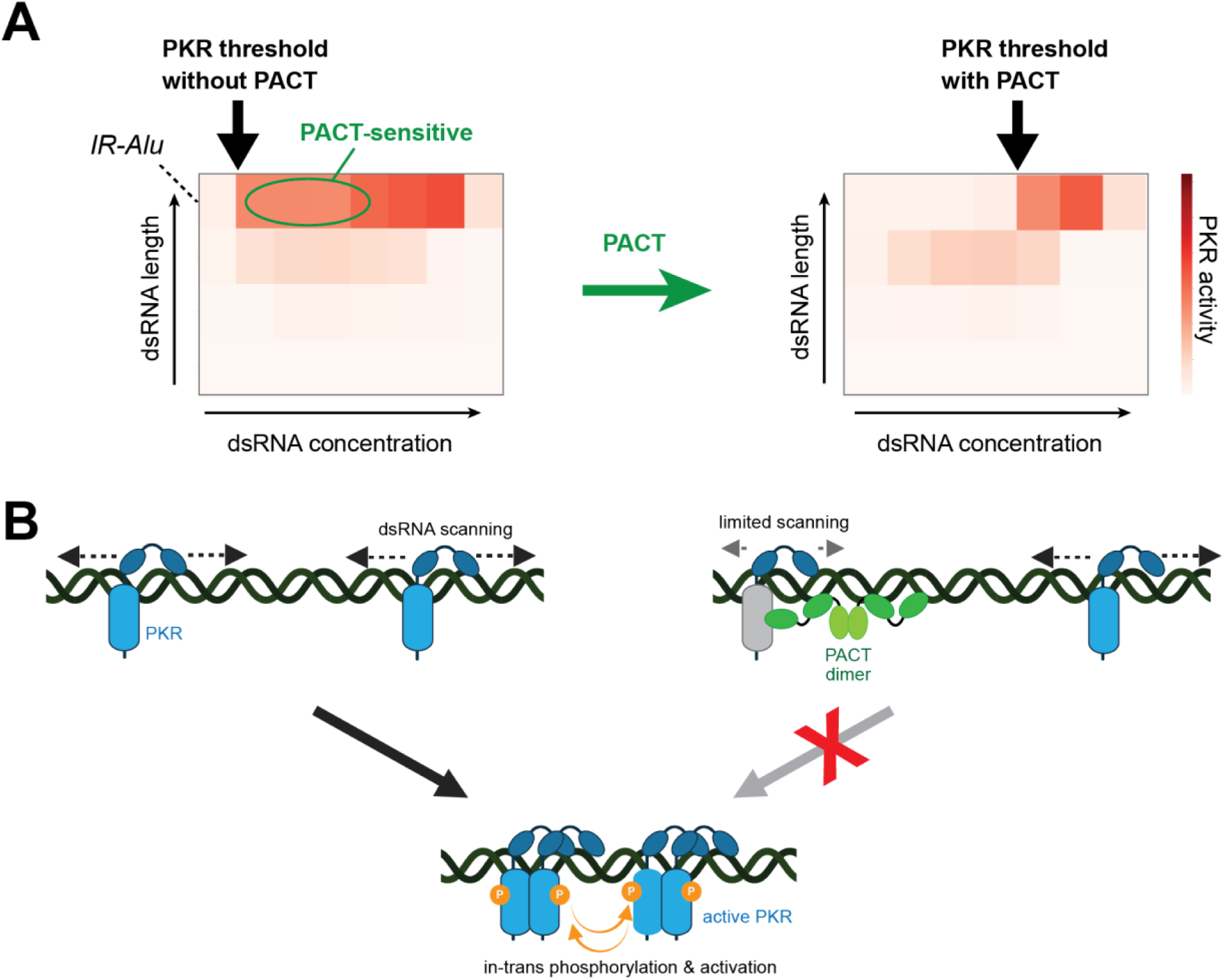
Model for PKR activation and PACT-mediated inhibition. (A) Heatmaps showing dsRNA length and concentration dependance for PKR activity in the absence and presence of PACT. PACT restricts PKR selectively within a specific range of the dsRNA length-concentration space, resulting in the selective elevation of PKR’s activity threshold for longer dsRNAs (such as IR-Alus). Notably, PACT has minimal impact on PKR when stimulated with high concentrations of long dsRNA, suggesting that PACT acts as a homeostatic regulator without compromising PKR’s inherent activity. (B) Model of PKR activation by dsRNA and its inhibition by PACT. PKR activation involves dsRNA scanning, molecular collisions and subsequent trans-autophosphorylation on long dsRNA. PACT also scans along dsRNA, restricting PKR’s movement, preventing molecular collision between PKR molecules. This inhibition requires more than PACT’s dsRNA-binding activity; it also involves a direct, albeit weak, interaction with PKR’s kinase domain, facilitated when both PKR and PACT are on the same dsRNA. As a result, PACT’s inhibition of PKR is more robust when dsRNA is longer and less abundant––conditions that promote co-occupancy–– explaining how PACT modulates its inhibitory function based on dsRNA length and concentration.

Our results also suggest that PACT’s inhibition involves more than simple dsRNA binding; it likely includes a direct interaction between PACT and the kinase domain of PKR. This interaction appeared weak in the absence of dsRNA, and would likely occur only when both PKR and PACT are bound to the same dsRNA. Although the low affinity of this interaction has made detailed characterization difficult, the phosphorylation of PACT by PKR, which requires direct contact, supports its existence. The observed correlation between PACT’s phosphorylation level and its ability to inhibit PKR further suggests a mechanistic link between the two processes. Moreover, mutations in PACT at putative interface regions, away from the dsRNA-binding domains, affect PKR activity both in vitro and in cells. These interactions may help explain why PACT specifically modulates PKR function with minimal effect on other dsRNA-binding proteins, such as ADAR1, OASes, and RLRs, though alternative explanations cannot be ruled out. Low-affinity interactions like these, which are often challenging to study, may be more common in nature than appreciated and could play important roles in mediating complex biological processes.

In summary, PACT’s ability to inhibit PKR in a dsRNA length- and concentration-dependent manner contrasts with other PKR inhibitors, such as viral inhibitors (e.g., vaccinia viral protein K3L), which use RNA-independent high-affinity interactions to globally inhibit PKR^56,57^. Our work thus reveals a new mechanism by which PKR and PACT function together to maintain immune homeostasis amid fluctuating levels of endogenous dsRNA.

## METHODS

### Material Preparation

#### Cell lines and culture conditions

All cancer cell lines were grown and maintained in RPMI media supplemented with 10% fetal bovine serum (FBS), and 1% penicillin, streptomycin, and L-glutamine except for NCI-H2023 which was maintained in DMEM/F12 media. The following cancer cell lines were obtained from American Type Culture Collection (ATCC): ZR-75-1, HCC1428, NCI-H1437, NCI-H2023, NCI-H1650, HCC1806, MDA-MB-157, NCI-H2286, NCI-H727 and SW1271. Prior to shipping each cell line, ATCC performs cell line authentication. All cell lines were tested for mycoplasma regularly.

#### Plasmids

Bacterial expression plasmids for PACT and PKR were made in pET47b and pET29b respectively. PACT wild type was inserted between BamHI and SalI sites. PKR was inserted between NdeI and KpnI sites. Point mutants and truncation mutants for PACT: ΔD1 (residues 124-313), ΔD2 (Δresidues 110-198), ΔD3 (residues 1-196), D3 (residues 235-313), and for PKR: ΔDRBDs (residues 170-551) and all point mutations were generated by site-directed mutagenesis. PACT DRBD3 (residues 235-313) was cloned in pET47b between BamHI and SalI sites. TRBP wild type was inserted in pET28a between NheI and HindIII sites and ADAR1 DRBDs (residues 496-800) were inserted in pET47b between BamHI and HindIII sites. RIG-I ΔCARDs and MDA5ΔCARDs were used from previous study^48,58^. For cellular studies, entry clones for *PACT* (wildtype, C77R, K173E, K187E, K191E, and 4KE) and *PKR K296R* were generated through PCR-amplification of the respective *PACT* and *PKR K296R* open reading frames (ORF) described above. These entry clones were sub-cloned into a Gateway donor vector. Overlap PCR was performed to introduce (1) silent mutations into the protospacer adjacent motif (PAM) sequences targeted by PACT sgRNA 1 for *PACT* clones or PKR sgRNA 1 for *PKR K296R* to render these constructs resistant to CRISPR-Cas9 editing by these sgRNAs; and (2) FLAG tags to the C terminus of *PACT* clones or a hemagglutinin tag to the C terminus of *PKR K296R*. The individual *PACT* constructs and a *GFP* control construct were sub-cloned into the pLEX306 lentiviral expression vector (Addgene) under the control of a human PGK promoter. The *PKR K296R* construct was sub-cloned into the pLEX307 lentiviral expression vector (Addgene) under the control of an EF-1α promoter. Each expression vector was then transfected into HEK 293T cells to generate lentivirus. Lentiviral transduction of target cell lines was performed as described below. For the *PACT* clones (wildtype, C77R, K173E, K187E, K191E, and 4KE), increasing volumes of lentivirus was used to transduce target cell lines to achieve a gradient of PACT protein expression in target cell lines.

#### Proteins

All PACT constructs (except 4KE, D3), TRBP and ADAR1 DRBDs (residue 496-800) were expressed and purified from *E. coli* BL21(DE3) using a combination of Ni-NTA, heparin affinity and size-exclusion chromatography. Cells were lyzed using high-pressure homogenization with Emulsiflex C3 (Avestin) and centrifuged. The supernatant was loaded onto Ni-NTA agarose beads and washed with 50 column volumes (CV) of high salt buffer (2M NaCl) to get rid of bound RNA contaminants before elution in 50 mM sodium phosphate pH 7.5, 1M NaCl, 10% glycerol, 300 mM imidazole. The eluate was further treated with HRV-3C protease and benzonase overnight to remove His-tag and contaminating RNAs respectively. After tag removal, the protein was subjected to heparin affinity chromatography followed by size-exclusion chromatography (SEC) in 50 mM HEPES pH 7.5, 250 mM NaCl and 2 mM DTT to further purify proteins and separate from benzonase. Removal of benzonase was confirmed by the stability of dsRNA co-incubated with purified protein at 37°C for 2 hr. For 4KE and D3 mutants, benzonase treatment and heparin affinity chromatography steps were skipped since they were RNA-binding defective.

PKR constructs were co-expressed and purified from *E. coli* Rosetta2 (DE3) (Novagen) using a combination of Ni-NTA and size-exclusion chromatography. NiNTA was done in the same buffer as PACT except the wash was done with 100 CV high salt buffer (50 mM Tris 8.0, 1M NaCl, 5mM β-mercaptoethanol (BME). The eluate from NiNTA was subjected to SEC in a 16/600 Superdex 200 column in 50 mM Tris pH 7.5, 300 mM NaCl, 5 mM BME. The eluate was sequentially treated with lambda protein phosphatase (PPase) (NEB) and benzonase (EMD Millipore) in the presence of 1 mM MnCl_2_ and 1 mM MgCl_2_ respectively. PKR was subjected to another round of SEC in 50 mM Tris pH 7.5, 100 mM NaCl and 5 mM BME to get rid of any contaminant PPase or benzonase. Note, that PKR wild type constructs were co-expressed with PPase expressing pET21b for purification. For K296R mutants, both PPase co-expression and post-SEC PPase treatment were skipped since the mutation is catalytically inactive. RIG-I ΔCARDs and MDA5 ΔCARDs were purified as reported previously^48,58^.

#### RNAs

All dsRNAs used in this study were generated by *in vitro* T7 transcription as reported previously^51^. The templates for RNA synthesis were made by PCR amplification. The sequences of 42, 62, 112 and 512 bp dsRNAs were taken from the first 30, 50, 100 and 500 bp of the MDA5 gene, respectively, flanked by 5’-gggaga and 5’-tctccc (See Table S3). The two complementary RNA strands were co-transcribed, and the duplex was purified using 8.5% acrylamide gel electrophoresis. RNA was gel-extracted using the Elutrap electroelution kit (Whatman), ethanol precipitated, and stored in 20 mM HEPES pH 7.0. For ^32^P labeled RNA, *in vitro* transcription was done in the presence of trace amounts of [α-^32^P]-UTP followed by RNA clean-up using RNA Clean & Concentrator Kit (Zymo research). The RNA concentration was estimated using Qubit Fluorometers (Thermo).

### CRISPR-Cas9 gene knockout

Single guide RNA sequences were designed using the sgRNA Designer tool (The Broad Institute Genomics Perturbation Platform) (http://portals. broadinstitute.org/gpp/public/analysis-tools/sgrna-design). sgRNA sequences are displayed in Supplementary Table S2. sgRNAs were cloned into the Cas9-expressing lentiviral vector lentiCRISPRv2 (Addgene) or a modified lentiCRISPRv2 construct that expresses two different sgRNAs under the control of separate human and mouse U6 promoters. Individual lentiCRISPRv2 vectors were introduced along with packaging vectors into HEK 293T cells via calcium phosphate transfection according to the manufacturer’s protocol (Clontech). Lentivirus was harvested at 48 and 72 h after transfection in RPMI media supplemented with 10% FBS and filtered with 45 μm filters before transduction of target cancer cell lines using centrifugation at 1000*g* for 2 hours in the presence of 8 mg/mL of polybrene (Santa Cruz Biotechnology). Transduced cell lines were selected in 2 μg/mL puromycin and/or 10 μg/mL blasticidin for at least 5 days prior to use in assays. Protein lysates were collected from the transduced cells and protein levels of the targeted gene(s) were assessed by immunoblotting.

### Cell viability assays

Cell counting was performed using a Coulter Particle Counter (Beckman-Coulter). For ATP bioluminescence experiments, cells were plated at a density of 3,000 cells per well in 96-well assay plates (Corning). ATP bioluminescence was assessed at 13 days after gene knockout with the CellTiter-Glo Luminescent Cell Viability Assay (Promega). For crystal violet staining assays, cells were plated at a density of 10,000 to 20,000 cells per well in 12 well tissue culture plates. Once the control cells grew to near confluency, each well was washed twice with ice-cold PBS, fixed on ice with ice cold methanol for 10 min, stained with 0.5% crystal violet solution (made with 25% methanol) for 10 min at room temperature, and washed at least four times with water. All cell viability assays were performed in triplicate.

### Antibodies and immunoblotting

Cells were lysed in RIPA lysis buffer (Thermo Fisher Scientific) supplemented with 1× protease and phosphatase inhibitor cocktails (Roche). Protein concentrations were obtained using the BCA Protein Assay Kit (Pierce) and 6X Laemmeli SDS sample buffer (Thermo Fisher Scientific) was added to protein extracts. Protein concentrations were normalized between all samples within an experiment and boiled above 95°C for 10 min. Proteins were resolved on 4-12% Bis-Tris gradient gels, transferred to nitrocellulose membranes, and immunoblotting with primary and secondary antibodies was performed according to standard procedures. All primary antibodies were used at a dilution of 1:1,000 except anti-β-actin and anti-vinculin, which were used at a dilution of 1:10,000. Secondary antibodies from LI-COR Biosciences were used at a dilution of 1:10,000. Selected immunoblots were stripped with Restore PLUS Western Blot Stripping buffer (Thermo Fisher Scientific) prior to repeat immunoblotting. Immunoblots were imaged using the LI-COR digital imaging system. All immunoblots were cropped to optimize clarity and presentation.

### *In vitro* PKR kinase assay

For RNA-dependent PKR kinase assays, 100 nM PKR was mixed with the mentioned concentration of RNA, 2 mM ATP and 1μCi [γ^32^P]-ATP in 20 mM HEPES pH 7.5, 100 mM NaCl, 1.5 mM MgCl_2_, 2 mM DTT in the presence of mentioned concentration of PACT and the reaction incubated at 30°C for 30 min. For RNA-dependent PKR kinase assays, 1 to 1.5μM PKR was first pre-treated with Benzonase (Millipore) in the presence of 2 mM MgCl_2_ at room temperature for 15 min. For the experiments involving PACT, PACT was also pre-treated with Benzonase (Millipore) in the presence of 2 mM MgCl_2_ at room temperature for 15 min. Both PACT and PKR, after benzonase treatment were mixed together and incubated at room temperature for 15 min before adding 2 mM ATP and 1μCi [γ^32^P]-ATP in the final buffer comprising of 20 mM HEPES pH 7.5, 150 mM NaCl, 1.5 mM MgCl_2_, 2 mM DTT and incubating at 37°C for 10 min. After the reaction incubation time, the samples were quenched in SDS gel loading buffer and subjected to SDS-PAGE. The gels were exposed to imaging plate from Cytiva and Phosphor imaging was done using Amersham Typhoon Imager (GE). Quantification was done using ImageJ.

### fCLIP-seq

The procedure was adopted from the previous study^3^. Cells were crosslinked with 0.1% paraformaldehyde for 10 min at RT followed by quenching with 150 mM glycine for 10 min at RT. The crosslinked cells were then resuspended in 20mM Tris pH 7.5, 15mM NaCl, 10mM EDTA, 0.5% NP-40, 0.1% Triton X-100, 0.1% sodium deoxycholate, 1x mammalian protease inhibitor cocktail (G Biosciences), 40U/ml RNase inhibitor (NEB) and 40U/ml DNase I (NEB) and incubated on ice for 10 min. The cells were lyzed by sonication using Covaris E220 focused-ultrasonicator (PIP: 140, DF: 5, Cycles/Burst: 200, Time: 5 min). The lysate was centrifuged at 18,000g for 30 min and the supernatant was added to pre-equilibrated protein G Dyanbeads (Invitrogen) along with Anti-HA (CST #3724), Anti-PKR (CST #12297) or Anti-Flag (Millipore Sigma #F1804) antibody. The mixture was incubated at 4°C for 2 h before washing beads with 0.1% SDS, 0.5% NP-40, 0.5% sodium deoxycholate in 1X PBS and eluting in 200 mM Tris-HCl pH 7.5, 100 mM NaCl, 20 mM EDTA, 2% SDS, and 7 M Urea with constant shaking at 950 rpm at 25°C for 1-2 h. The eluted protein:RNA complex was then de-crosslinked overnight at 65°C with constant shaking at 1000 rpm in the presence of 2.7 U/ml proteinase K (NEB) to digest the proteins. The RNA was then extracted by phenol:chloroform extraction followed by isopropanol precipitation. The extracted RNA was further treated with DNase I (NEB) for 30 min at 37°C to completely get rid of any DNA contamination. The purified RNA used for cDNA library preparation using SMARTer Stranded Total RNA-Seq Kit v3 - Pico Input Mammalian according to manufacturer’s instructions. The cDNA library was sequenced using Illumina Partiallane NovaSeq platform.

### RT-qPCR

Total RNAs were extracted using Directzol RNA miniprep kit (Zymo research) and cDNA was synthesized using High Capacity cDNA reverse transcription kit (Applied Biosystems) according to the manufacture’s instruction. Real-time PCR was performed using a set of gene specific primers, SYBR Green Master Mix (Applied Biosystems), and the CFX96 Real-Time PCR Systems (Biorad).

### Native gel-shift assay

For quantitative analysis, 0.2 ng/μl ^32^P labelled 112 bp was incubated with 0, 3.9, 7.8, 15.6, 31.3, 62.5, 125, 250, 500, 1000, 2000, 4000 nM PACT proteins or TRBP, RIG-I ΔCARDs, ADAR1 DRBDs in 20 mM HEPES pH 7.5, 150 mM NaCl, 2 mM DTT and incubated at RT for 15 min. The samples were then subjected to Bis-Tris native PAGE (Life Technologies). The gels were exposed to imaging plate from Cytiva and Phosphor imaging was done using Amersham Typhoon Imager (GE) followed by quantification in ImageJ. The quantification was done as previously reported^51^. Briefly, the bound fraction was calculated using the equation:

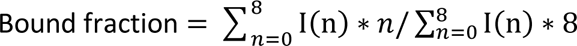

where I(n) refers to the intensity of the nth complex band. Free dsRNA is referred to by the 0th complex band. The underlying assumption is that 112 bp dsRNA has eight binding sites and the nth complex band corresponds to dsRNA with n sites occupied.

For non-quantitative analysis 25 ng/μl RNA (42 or 62 bp) was incubated with 0, 78, 156, 312.5, 625, 1250, 2500nM nM PKR at RT for 15 min before subjecting to native PAGE. The gel was stained with Sybr Gold (Thermo) and imaged using iBright FL1000 (Invitrogen).

### Multi-Angle Light Scattering (MALS)

PKR K296R mutant (0.7 mg/ml) was mixed with 50 ng/μl RNA (42 or 62 bp) in 20 mM HEPES pH 7.5, 150 mM NaCl, 2 mM DTT and incubated at RT for 30 min. The mix was then filtered using a 0.22 μ filter and then loaded onto Superdex 200 Increase 10/300 column (Cytiva) attached to MiniDAWN detector (Wyatt Technology) in 20 mM HEPES pH 7.5, 150 mM NaCl, 2 mM DTT. The data were analyzed using ASTRA7.3.1 software (Wyatt Technology).

### Electron Microscopy

PKR, PACT or MDA5ΔCARDs (0.3 μM) was incubated with 512 bp dsRNA (0.6 ng/μl) in 20 mM HEPES pH 7.5, 150 mM NaCl, 2 mM DTT and incubated at RT for 15 min. The samples were, then, adsorbed to carbon-coated grids (Electron Microscopy Sciences) and stained with 0.75% uranyl formate as described^59^. Images were collected using the JEM-1400 transmission electron microscope (JEOL) at a magnification of 30,000x.

### GST pulldown

GST-tagged PKR (K296R) (0.7 μM) and PACT (5 μM) were individually benzonase-treated for 15 min at RT in the presence of 1 mM MgCl_2_. Following the benzonase treatment, both proteins were mixed together and incubated for another 15 min at RT. The protein mix was then incubated at 37°C for 10 min in 20 mM HEPES pH 7.5, 150 mM NaCl, 2 mM DTT before adding to Glutathione Sepharose 4B beads (Cytiva) pre-equilibrated with 50 mM Tris pH 7.5, 250 mM NaCl and incubated at 4°C overnight with gentle rocking. The beads were then washed 3 times with 50 mM Tris pH 7.5, 250 mM NaCl before eluting with 50 mM reduced glutathione in 50 mM Tris, 150 mM NaCl. The samples were subjected to SDS-PAGE and analyzed by Western blot. The method for PKR pulldown by PACT D1D2-GST was the same as above except that 0.45mg/ml PACT D1D2-GST and 0.35 mg/ml of PKR (K296R), full-length or ΔDRBD were used.

### Microscale Thermophoresis

PACT ΔD3 was purified using the method described earlier but a fluorescent label was added prior to the heparin chromatography step. This was achieved by incubating the protein with 1 mg/ml *S. aureus* sortase A (a gift from H. Ploegh, MIT)^60^ and 100 μM peptide (LPETGG) conjugated with FAM (Anaspec) for 2 h at RT protected from light. The FAM-labeled PACT ΔD3 GST-tagged PKR (K296R) were pre-treated with benzonase at RT for 15 min in the presence of 1 mM MgCl_2_ to remove any RNA contaminants. Post-benzonase treatment FAM-labeled PACT ΔD3 (1 μM) was incubated with 16 different concentrations of GST-tagged PKR (K296R) (5 μM to 0.000458 μM) in 20 mM HEPES pH 7.5, 150 mM NaCl, 2 mM DTT, 0.05% Tween20 for 30 min at RT in the dark. Each titration mix was loaded in a separate capillary and the MST scan was recorded on a Monolith NT.115pico (NanoTemper Technologies) using the nano Blue detector at the Center for Macromolecular Interactions, Harvard Medical School. Raw fluorescence for each capillary was scanned before and after the MST trace to ensure that the samples did not aggregate during the course of MST experiment. The data from 3 independent experiments were analyzed using MO.Affinity Analysis v2.3.

### Bioanalyzer analysis

Total RNA from 5 different cell lines (HCC1806, NCI-H727, NCI-H2286, NCI-H2023, NCI-H1437) was isolated on day 6 after control or PACT knockout. For RNA isolation, TRIzol reagent (Thermo) was added to the cells after removing media and washing once with phosphate buffer saline (PBS). The TRIzol-mixed cell lysate was used for RNA extraction using Directzol RNA miniprep kit (Zymo research). The purified RNA was loaded on RNA nano chip using an Agilent Bioanalyzer.

### NMR

#### Sample preparation

PACT D3 triple labeled (^15^N, ^13^C, ∼85% ^2^H) sample was prepared by expressing the protein in *E. coli* BL21(DE3) grown in M9 minimal media supplemented with ^15^NH_4_Cl (99%), ^13^C D-glucose (99%) as the sole sources of N and C respectively. The minimal media was made in deuterium oxide instead of H_2_O in order to ensure ^2^H incorporation in the protein. The purification method was the same as described in previous section except that final SEC was done in 20 mM HEPES pH 7.5, 50 mM KCl, 2 mM DTT. For mixed isotope sample, 2 separate preparations were carried out: one with ^15^NH_4_Cl (99%) in deuterium oxide M9 minimal media for ^15^N, ^2^H-labeled protomer sample and the other with ^13^C D-glucose (99%) in regular M9 minimal media for ^13^C-labeled protomer. Both protomers were individually purified and then denatured by adding 6 volumes of 7M guanidinium hydrochloride along with 20 mM BME and shaking at 37°C for 1 h. The two denatured protomers were then mixed in equimolar ratio and dialyzed overnight in 50 mM HEPES pH 7.5, 150 mM NaCl, 2 mM DTT at 4°C using a 3 kDa MWCO. The refolded mixed isotope sample was concentrated and subjected to SEC in 20 mM HEPES pH 7.5, 50 mM KCl, 2 mM DTT.

#### Assignment of NMR Resonances and NOE Restraints

All NMR data were recorded at 30 °C (303 K) on Bruker spectrometers operating at ^1^H frequency of 900 MHz or 600 MHz and equipped with cryogenic probes. NMR data were processed using NMRPipe^61^ and spectra analysis was performed using XEASY^62^ and CcpNmr^63^. Triple resonance experiments were collected at ^1^H frequency of 600 MHz using a (^15^N, ^13^C, ∼85% ^2^H)-labeled sample. Sequence specific assignment of backbone chemical shifts was accomplished using three pairs of TROSY-enhanced triple resonance experiments^64,65^, including HNCA, HN(CO)CA, HN(CA)CO, HNCO, HNCACB, and HN(CO)CACB. The aliphatic and aromatic resonances of the protein sidechains were assigned using the 3D ^15^N-edited NOESY-TROSY-HSQC (τ_NOE_ = 80 ms) and 3D ^13^C-edited NOESY-HSQC (τ_NOE_ = 150 ms) spectra, recorded at ^1^H frequency of 900 MHz using a (^15^N, ^13^C)-labeled protein sample. These NOE spectra were also used for assigning intra-chain NOEs. For assigning inter-chain distance restraints, a mixed sample was prepared in which half of the protomers were ^15^N, ^2^H-labeled (0.4 mM) and the other half 15% ^13^C-labeled (0.4 mM). Recording a 3D ^15^N-edited NOESY-TROSY-HSQC (τ_NOE_ = 200 ms) on this sample allowed measurement of exclusive NOEs between the ^15^N-attached protons of one subunit and aliphatic protons of the neighboring subunits. The non-deuterated protein was 15% ^13^C-labeled for recording the ^1^H-^13^C HSQC spectrum as internal aliphatic proton chemical shift reference.

#### Structure calculation

Structure calculation was performed using the program XPLOR-NIH^66^, involving two steps: 1) obtaining the structure of a monomeric subunit and 2) determining the dimeric assembly that best satisfy the intermolecular NOE restraints. In the first step, we calculated the monomer structure using all intramolecular NOE restraints, hydrogen bond restraints, and backbone dihedral angles derived from ^1^H_N_, ^15^N, ^13^C_α_, ^13^C_β_, and ^13^C’ chemical shifts using the TALOS+ program^67^. For this step, we used a simulated annealing (SA) protocol in which the temperature in the bath was cooled from 2000 to 200 K with steps of 40 K. The distance restraints were enforced by flat-well harmonic potentials, with the force constant ramped from 2 to 30 kcal/mol Å^-2^ during annealing. Backbone dihedral angle restraints were taken from the ‘GOOD’ dihedral angles from TALOS+, all with a flat-well (±the corresponding uncertainties from TALOS+) harmonic potential with force constant ramped from 30 to 100 kcal/mol rad^-2^. A total of 50 structures were calculated and the lowest energy structure was selected as the monomer structure for subsequent dimer structure calculation. For dimer calculation, we generated two identical copies of the monomer structure from Step 1 and performed a similar SA protocol while applying all NMR restraints including the inter-chain distance restraints. In this protocol, the bath was cooled from 1000 to 200 K with steps of 20 K. The NMR restraints were enforced as in Step 1. A total of 75 structures were calculated and 15 lowest energy structures were selected as the final structural ensemble.

##### Single-molecule FRET measurements

For 3’-labeling with Cy3 or biotin, 112 nt RNA was oxidized with 0.1 M sodium meta-periodate (Pierce) overnight in 0.1 M NaOAc pH 5.4. The reaction was quenched with 250 mM KCl, buffer exchanged using Zaba desalting columns (Thermo Fischer) into 0.1 M NaOAc pH 5.4 and further incubated with 0.1 M Cy3-Mono-hydrazide (GE Healthcare) or EZ-Link Hydrazide-PEG4-Biotin (Pierce) for Cy3 or biotin labeling respectively for 4–6 hr at RT. The dsRNA sample was then prepared by annealing 3’-Cy3 forward strand (112 nt) and 3’-biotin labeled reverse complementary strand by mixing them in equimolar ratio to 95°C followed by slowly cooling down to 25°C at a ramp speed of −0.5°C/s. PKR K296R mutant or PACT was labeled with Cy5 using Sortase A as described above.

All the single molecule experiments were carried out using a home-built prism-type total internal reflection fluorescence microscope at room temperature (23.0 ± 1.0 °C). Cy3 labeled dsRNA was diluted to 200 pM with 20 µL of 10 mM pH 8.5 Tris-HCl and 50 mM NaCl (T50) and immobilized on a PEG-coated quartz slide pretreated with neutravidin (0.05 mg/ml). To minimize bleaching and blinking of dyes, an imaging buffer was prepared with reaction buffer (1mg/ml glucose oxidase, 0.8 % v/v dextrose, ∼ 2mM Trolox, 0.03 mg/ml catalase, 20mM HEPES pH 7.5 and 50mM NaCl). 532 nm solid-state diode laser was employed to excite Cy3. Fluorescent emissions of Cy3 and Cy5 were separated by a dichroic mirror (cutoff: 640 nm) and collected by electron-multiplying CCD (Andor). All data was recorded with a 33 ms time resolution by smCamera software and analyzed with Interactive Data Language. Single-molecule traces and histograms are further analyzed with scripts written in MATLAB. IDL and Matlab scripts were provided by Dr.Taekjip Ha’s lab. The duration time and sliding time were fitted using Origin 2018.

##### Data analysis for single-molecule FRET

The auto-correlation function was used to analyze the sliding time of PKR, PACT and PKR in the presence of PACT, as described in^54^. The equation of the auto-correlation function is as below:

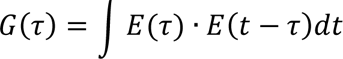

*E*(*τ*) represents FRET, and the FRET auto-correlation, *G*(*τ*), gave us an average sliding time because FRET changes in our system reflect the diffusion of a protein. More than 200 binding events with 3-5 movies were analyzed for each auto-correlation curve, and it was fitted with a single exponential decay to obtain an average sliding time of the protein. Duration times of the diffusive state were measured from individual traces, and were plotted by Prism10. To compare the movement range of PKR with and without PACT, we performed the histogram analysis of FRET states. Aggregate analysis of all traces was unable to distinguish between movements confined to low-, mid- or high-FRET state, vs. those that fluctuate between low and high states. Therefore, we limited our analysis to traces showing high FRET and compiled these traces to generate the FRET histogram for 50 nM PKR with and without PACT.

### Analysis of sequencing data

#### RNA-seq alignment and normalization

Pair-ended fastq samples were first trimmed by trimmomatic (v0.36)^68^, and mapped to *Homo sapiens* RNA45SN1 (NCBI Gene ID: 106631777) and RNA5S1 (NCBI Gene ID: 100169751). The leftover reads were then mapped to human genome (GRCh38.p14). Only primary alignments were kept by SAMtools (v1.9)^69^ flag -F 260. To adjust for different sequencing depths among samples, the bam files were first converted to bedgraphs by bedtools genomecov (v2.30.0)^70^, and the bedgraph coverage values were normalized to “as if one million total non-rRNA reads were mapped to human genome” by multiplying the coverage value with a factor of 10^6/(total non-rRNA mapped read counts). The normalized bedgraphs were named as “*PM.bedgraph” for individual groups in our GEO submission.

#### Bedgraph peak calling and AuC calculation

The normalized fCLIP bedgraphs were first subtracted with normalized input bedgraphs with macs2 (v2.1.1.20160309) bdgcmp ^71^, and the resultant bedgraphs were used for peak calling with macs2 bdgpeakcall ^71^. The individual AuCs were calculated from these peak regions using bedtools (v2.30.0) intersect with individual PM bedgraphs, and then calculating the sum of all overlapping bedgraph regions in the same peak.

#### Inverted Repeat calculation

Repeat regions in the human genome were acquired from Repbase (girinst.org/repbase). For all repeat regions of the same family, their sequences were searched for similarity by NCBI BLAST (blastn v2.15.0; reward=2, penalty=-3, gapopen=5, gapextend=2, word_size=11) with neighboring repeats up to 1.5 kb. If their whole sequences match over 80% percentage identity with plus/minus match, they were then identified as inverted repeats.

#### Deseq2 calculation of AuC values

The adjusted p-value and FC of AuC values in peak regions were calculated by Deseq2^72^ in R. Using peak regions as the index, the Deseq2 result datasheet was merged with other associated information (e.g. inverted repeats) using peak name as index by python pandas.

#### Github repository

In our Github repository, you will find detailed documentation of:

1. sbatch shell script to trim and map the fastq files using Harvard Medical School (HMS) O2 cloud computing platform
2. sbatch shell script to normalize bam files, convert to bedgraph, subtract input from fCLIP, call peak and individual AuC calculation
3. python script of calculating Inverted Repeat regions using NCBI BLAST engine

Github link: https://github.com/DylannnWX/Hurlab_PKR_manuscript/tree/main.

## Data and code availability

The NMR data in this study is deposited Biological Magnetic Resonance Bank with the accession code BMRB: 36671. The atomic coordinates have been deposited in the Protein Data Bank with accession code PDB: 8ZU6. The accession code for the fCLIP-seq data is NCBI GEO dataset: GSE269684.

## Supplementary Figures

**Figure S1.**
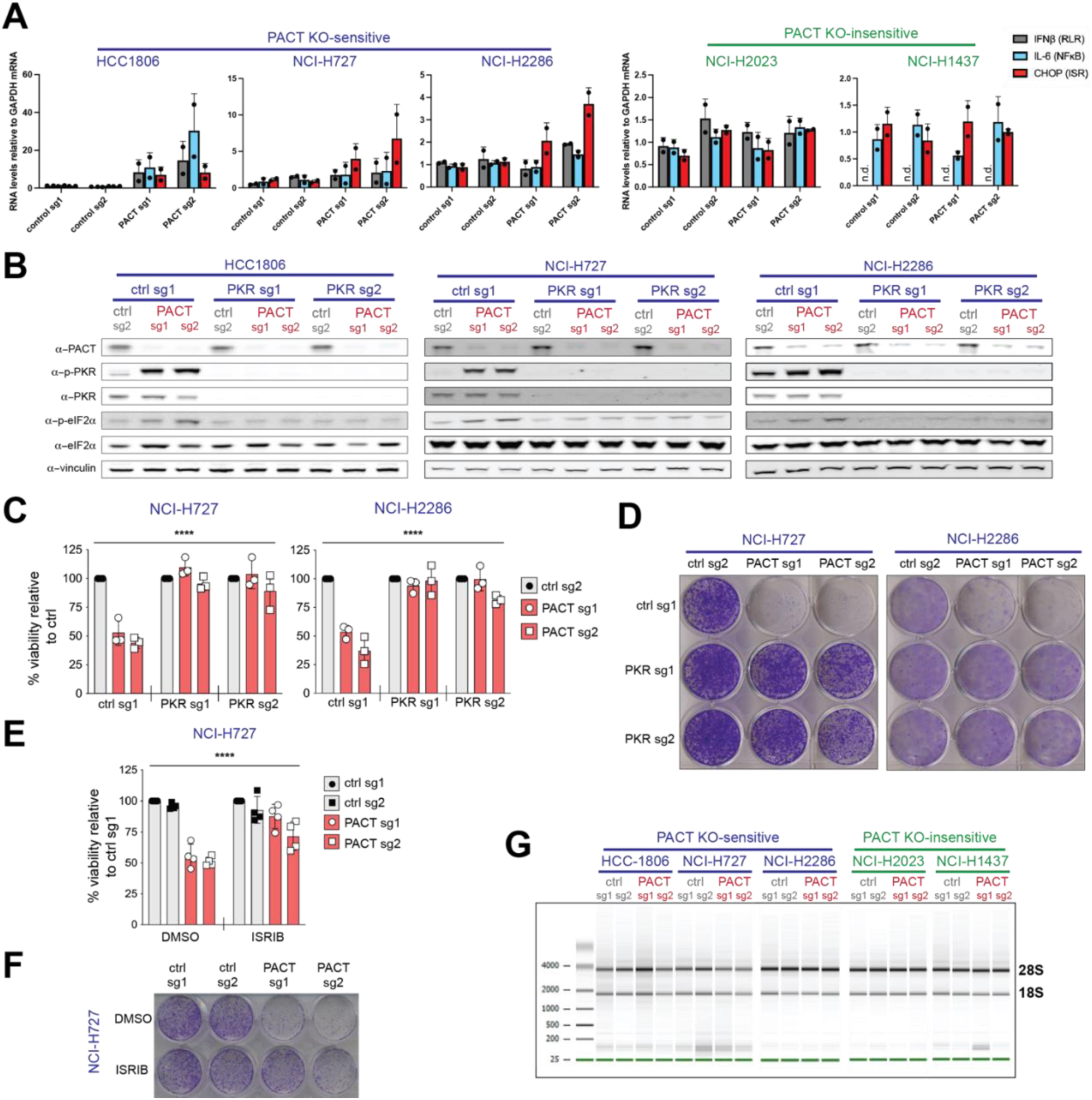
PACT restricts aberrant activation of PKR, thereby maintaining cellular homeostasis. (A) RT-qPCR analysis with RNAs extracted from PACT KO-sensitive (HCC1806, NCI-H727, NCI-H2286) and PACT KO-insensitive (NCI-H2023 and NCI-H1437) cells after control gene-KO and PACT-KO using 2 different guide RNAs. *IFNβ*, *IL-6* and *CHOP* mRNA levels relative to *GAPDH* were measured. The control genes are AAVS1 (sg1) and Chr2.2 (sg2). (B) Western blot analysis showing levels of PKR and eIF2α phosphorylation in HCC1806, NCI-H727, NCI-H2286 cells (PACT KO-sensitive cells) after knocking out indicated genes. Vinculin was used as a loading control. (C) ATP bioluminescence cell viability assay with NCI-H727 and NCI-H2286 cells after knocking out indicated genes. Values represent means of 3 biological repeats. *P*-values were based on two-way ANOVA test. (D) A representative crystal violet staining assay result for samples in (C). (E) ATP bioluminescence cell viability assay with NCI-H727 cells in control gene-KO vs. PACT-KO treated with 1μM ISRIB or DMSO. Values represent means of 4 technical repeats. *P*-values were based on two-way ANOVA test. (F) A representative crystal violet staining assay result from samples in (E). (G) Bioanalyzer run with total RNA extracted from PACT KO-sensitive (HCC1806, NCI-H727, NCI-H2286) and PACT KO-insensitive (NCI-H2023 and NCI-H1437) cells after knocking out indicated genes. **** *p*= <0.0001, *** *p* = <0.001, ** *p* = <0.01, * *p* = <0.05, not significant (ns) is for *p* > 0.05

**Figure S2.**
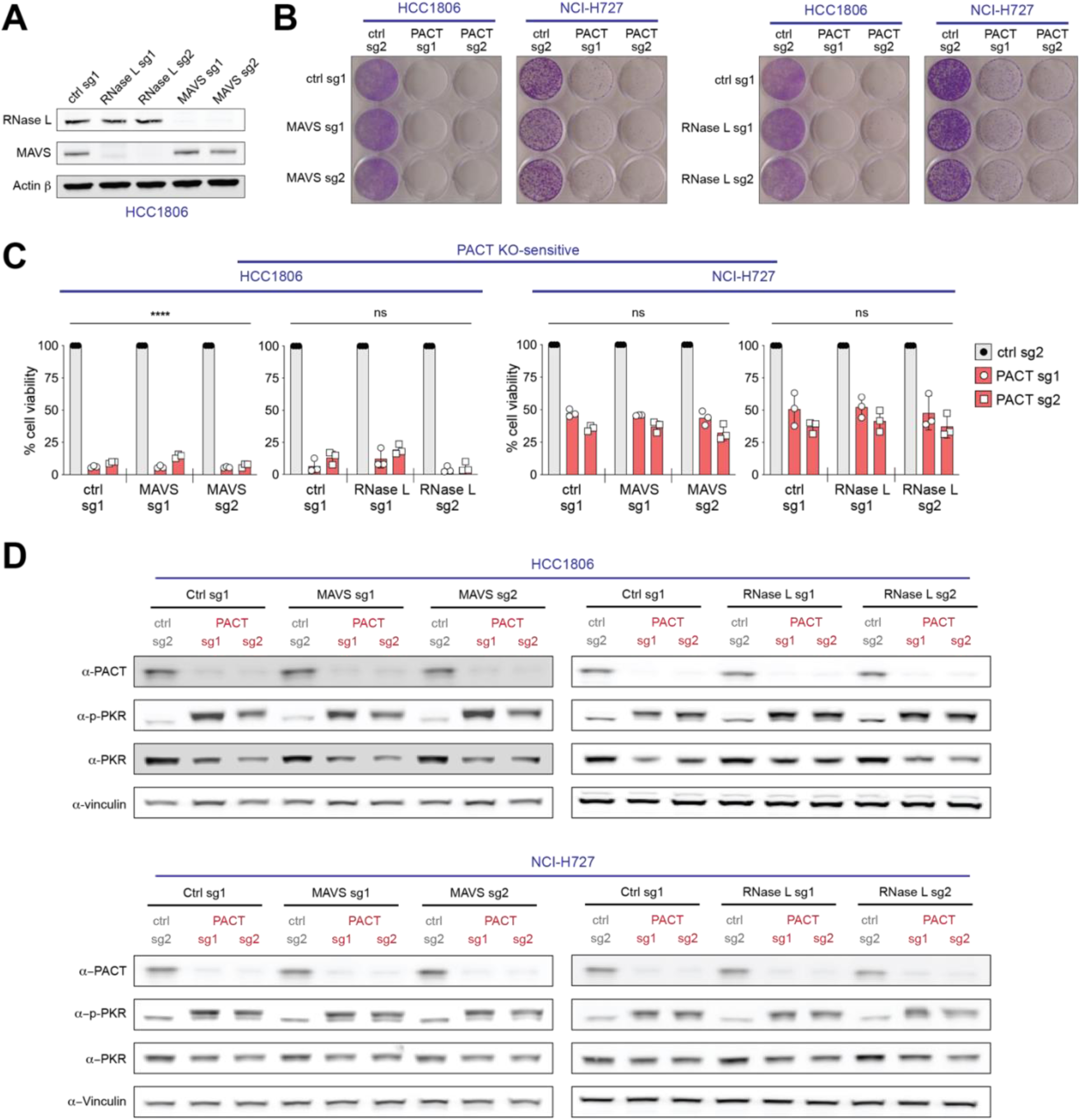
PACT has little role in RLR-MAVS and OAS-RNase L pathways. (A) Western blot analysis of HCC1806 cells after knocking out indicated genes. (B) Crystal violet staining assay showing cell viability after knocking out indicated genes in HCC1806 and NCI-H727 cells. (C) ATP bioluminescence assay showing cell viability after knocking out indicated genes in HCC1806 and NCI-H727 cells. Values represent means of 3 biological repeats. *P*-values were based on two-way ANOVA test. **** *p*= <0.0001, *** *p* = <0.001, ** *p* = <0.01, * *p* = <0.05, not significant (ns) is for *p* >0.05. (D) Western blot analysis with HCC1806 (top) and NCI-H727 (bottom) cells used in (B, C) showing PKR phosphorylation levels. Vinculin was used as loading control.

**Figure S3.**
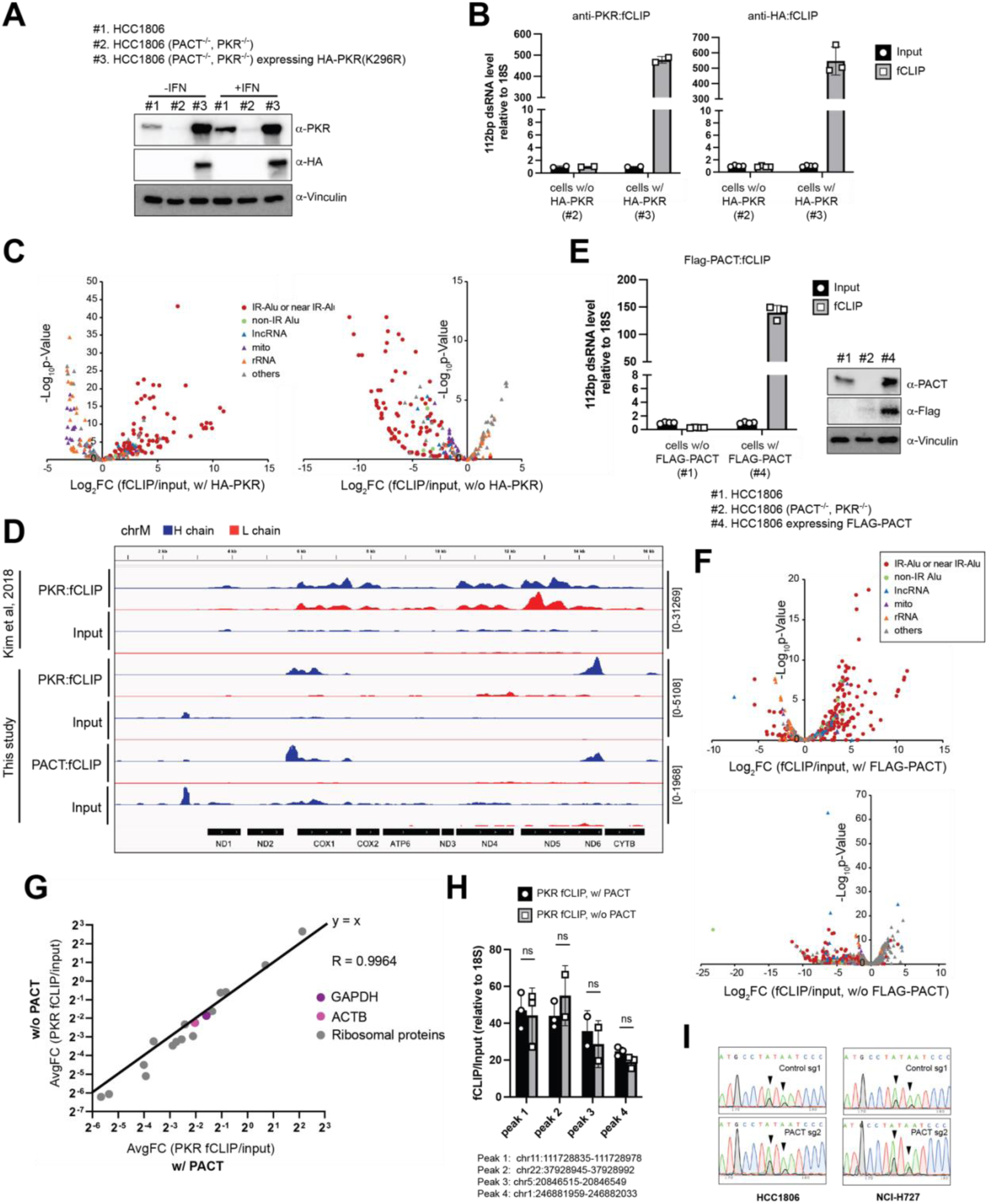
PACT does not sequester endogenous dsRNA ligands away from PKR. (A) Western blot analysis of PKR and PACT upon double knockout and complementation with catalytically dead PKR K296R mutant in HCC1806 cells in the presence and absence of IFNβ treatment (18 hr). (B) fCLIP-RT-qPCR showing enrichment of transfected 112bp dsRNA relative to 18S rRNA. HCC1806 cells expressing HA-PKR (#3 from A) and cells without HA-PKR (#2 from A) were compared. fCLIP was done using α-PKR or α-HA antibody. Values represent means of 2 and 4 technical replicates for α-PKR or α-HA antibodies respectively. (C) Volcano plots showing peaks enriched in fCLIP-seq using HCC1806 cells expressing HA-PKR (left, #3 cell from A) or not (right, #2 cells from A). fCLIP was done using α-HA antibody. Fold change (FC) of fCLIP over input was calculated. Values represent means of 3 biological replicates. (D) IGV snapshots showing the mitochondrial RNA peaks from this study (PKR and PACT fCLIP) compared to the previously reported PKR fCLIP (Kim *et al*, 2018). All RNA-seq data are strand-specific, allowing separate mapping of the H and L chains. While Kim *et al*. reported a significant accumulation of the L chain, our analysis showed relatively low levels of the L chain compared to the H chain. This discrepancy may reflect cell type-dependent variability in mitochondrial RNA synthesis/metabolism (Hela in Kim *et al* vs. HCC1806 in this study) or the effect of cell-cycle arrest treatment, under which the previous study was performed. (E) fCLIP-RT-qPCR showing enrichment of transfected 112bp dsRNA relative to 18S rRNA. HCC1806 cells expressing FLAG-PACT (#4 from E) and cells without FLAG-PACT (#1 from E) were compared. fCLIP was done using anti-Flag antibody. Values represent means of 4 technical replicates. Right: WB showing the levels of PACT in individual cell lines. (F) Volcano plots showing peaks enriched in fCLIP-seq using HCC1806 cells expressing FLAG-PACT (top, #4 cell from A) or not (bottom, #1 cells from E). fCLIP was done using α-FLAG antibody. Fold change (FC) of fCLIP over input was calculated. Values represent means of 2 biological replicates. (G) Validation of the normalization strategies for fCLIP-seq in Figure 2. We examined 17 distinct, unrelated housekeeping genes—including ACTB, GAPDH, and various ribosomal protein genes— from HA-PKR fCLIP with and without PACT. While these genes do not harbor fCLIP peaks that meet the filtering criteria in Figure 2C, they still have read counts from HA-PKR fCLIP-seq, allowing us to assess the normalization strategies. The values were plotted using the same method as in Figure 2F. The results showed nearly identical PKR fCLIP-to-input fold change of house keeping genes in the presence and absence of PACT (with the correlation coefficient R of 0.9964), suggesting equivalent background levels. (H) Re-examination of 4 peaks selected from (Figure 2F) by fCLIP RT-qPCR. Experiments were performed as in Figure 2F. Values were obtained by first normalizing indicated RNA levels to the internal control 18S rRNA, and then calculating the ratio of fCLIP to input for the normalized RNA levels. The results represent mean ± SD from 3 independent experiments. *P*-values were calculated by two-tailed t-test. (ns, not significant; *P*>0.05). (I) Representative sequencing chromatograms showing A-to-I editing sites in *PHAX* mRNA 3’UTR Alu sequence in control vs. PACT-KO in HCC1806 and NCI-H727 cells.

**Figure S4.**
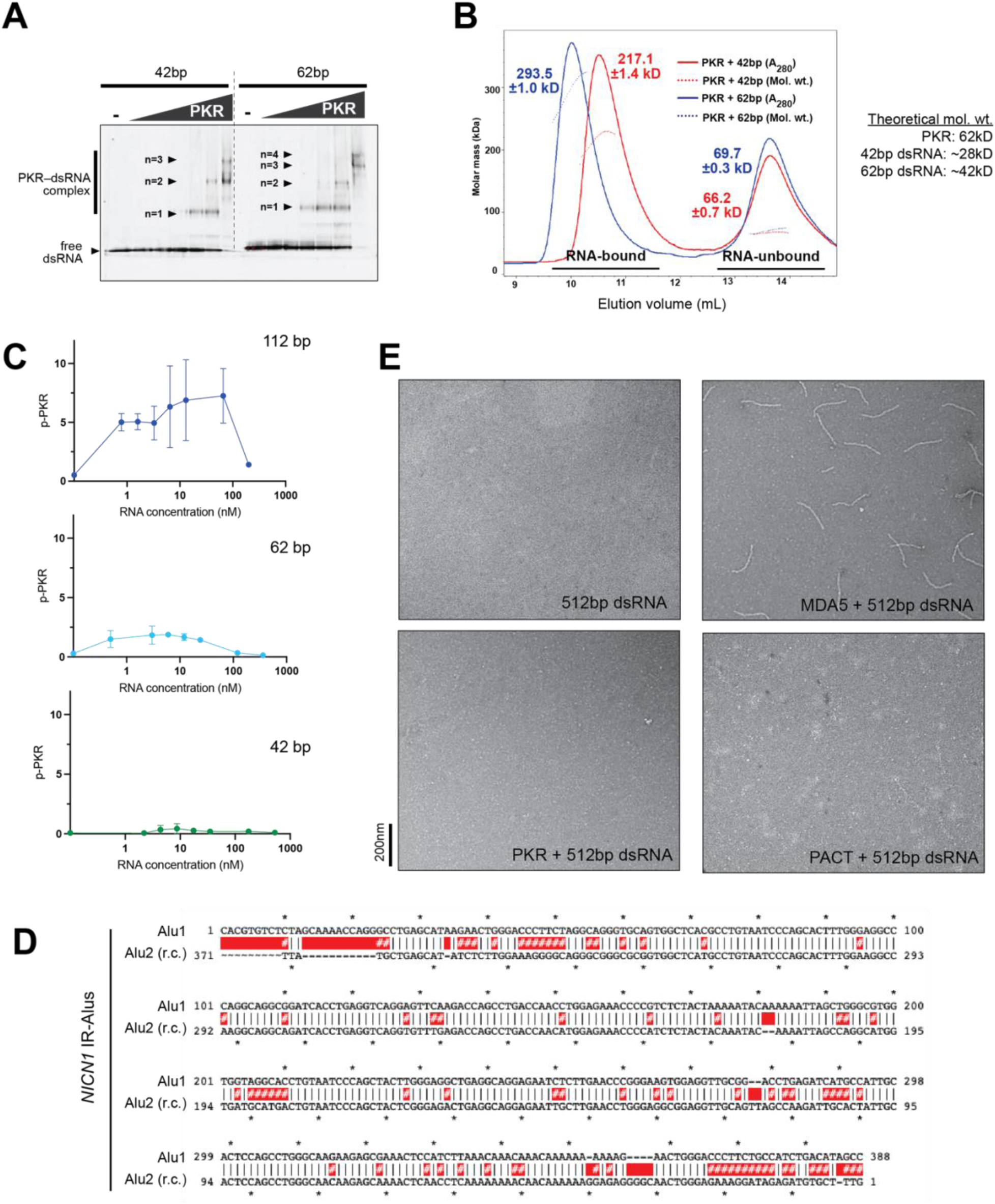
PKR can measure dsRNA length beyond 30 bp without forming filaments. (A) Native gel-shift assay showing binding of increasing concentrations (0, 78.1, 156.25, 312.5 625, 1250, 2500 nM) of PKR to 42 and 62 bp dsRNA (25 ng/μl). Gels were stained with Sybr gold. The number of complex bands suggests that 42 and 62 bp dsRNA can be occupied by up to 3 and 4 PKR molecules, respectively. (B) SEC-MALS analysis with PKR bound to 42 bp (red) and 62 bp (blue) dsRNA. Catalytic dead K296R mutant of PKR was used. Estimated molecular weights of the complexes are consistent with the notion that 42 bp and 62 bp dsRNAs are bound by 3 and 4 PKR molecules, respectively. (C) *In vitro* PKR kinase assay results in Figure 3D, but shown as a function of molar concentrations of RNA, rather than mass concentration. (D) Complementarity of *NICN1* IR-Alus, represented by alignment between the first Alu in the pair (Alu1) and reverse complement (r.c.) of the second Alu (Alu2). Vertical lines indicate complementarity, while red highlights indicate mismatches (#) or bulges (no symbol). (E) Negative-stain EM of 512 bp dsRNA with and without 300 nM MDA5ΔCARDs, PKR, PACT. While MDA5 forms filaments along the length of dsRNA, PKR and PACT do not form similar assemblies.

**Figure S5.**
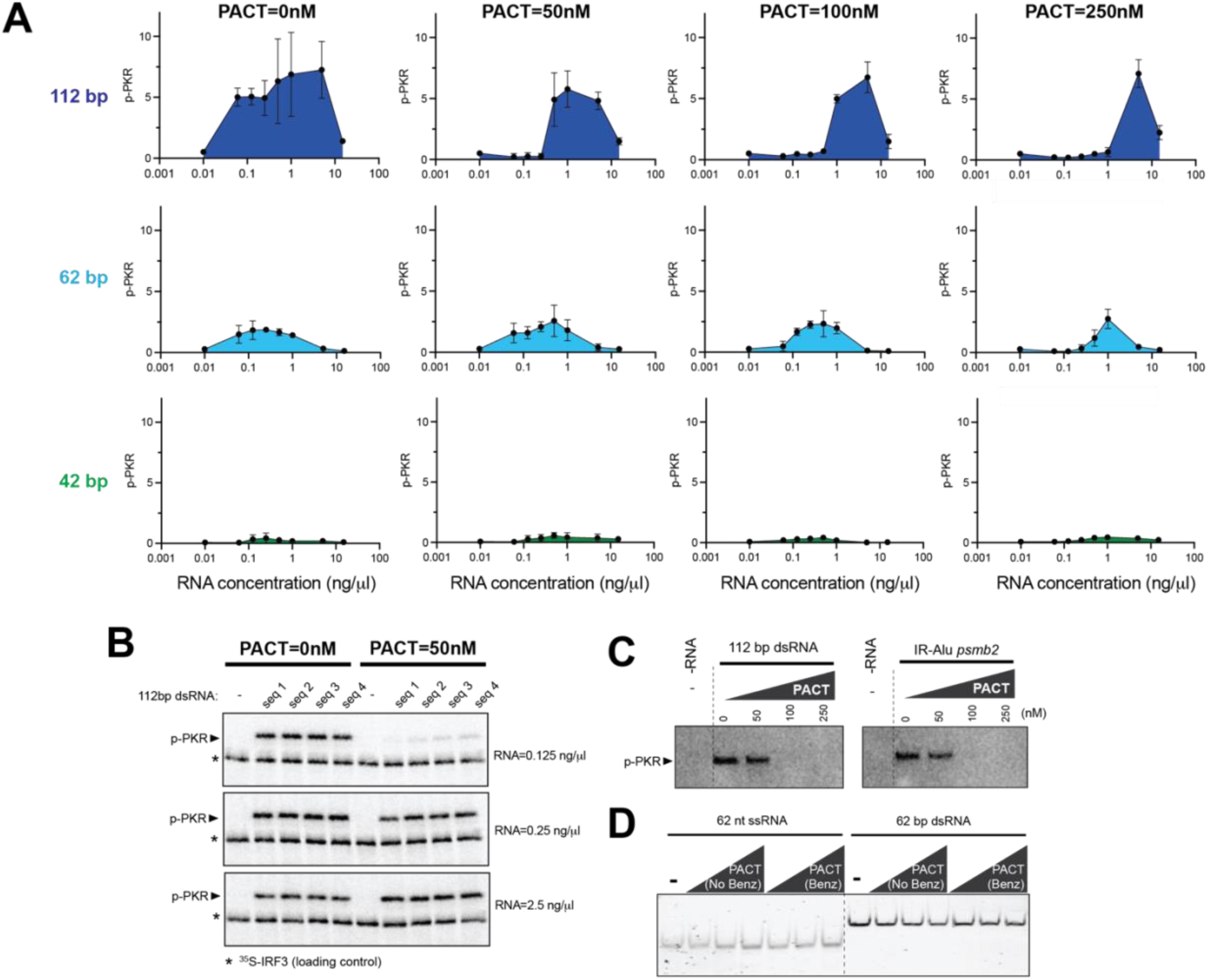
PACT’s ability to inhibit PKR is also dependent on dsRNA length. (A) *In vitro* PKR kinase assay results from Figure 4A. Values represent means (±SD) of 3 biological repeats. (B) *In vitro* PKR kinase assay using 112 bp dsRNA with 4 different sequences, seq 1 (GC%=52.7), seq 2 (GC%=50.1), seq 3 (GC%=42.9) and seq 4 (GC%=58.0) at 0.125. 0.25 and 2.5 ng/μl in the presence of 0, 50 nM PACT. See table S3 for RNA sequences. Results without PACT was reproduced from Figure 3A. (C) *In vitro* PKR kinase assay using 112 bp dsRNA or *PSMB2* IR-Alus (0.25 ng/μl) in the presence of 0, 50, 100, 250 nM PACT. (D) Benzonase contamination test by incubating 62 nt ssRNA or 62 bp dsRNA with increasing concentration (50, 100, 200nM) of PACT that was either purified without benzonase treatment (No Benz) or with benzonase treatment (Benz). The incubation was done for 2 h at 37°C in the presence of 1.5 mM MgCl_2_.

**Figure S6.**
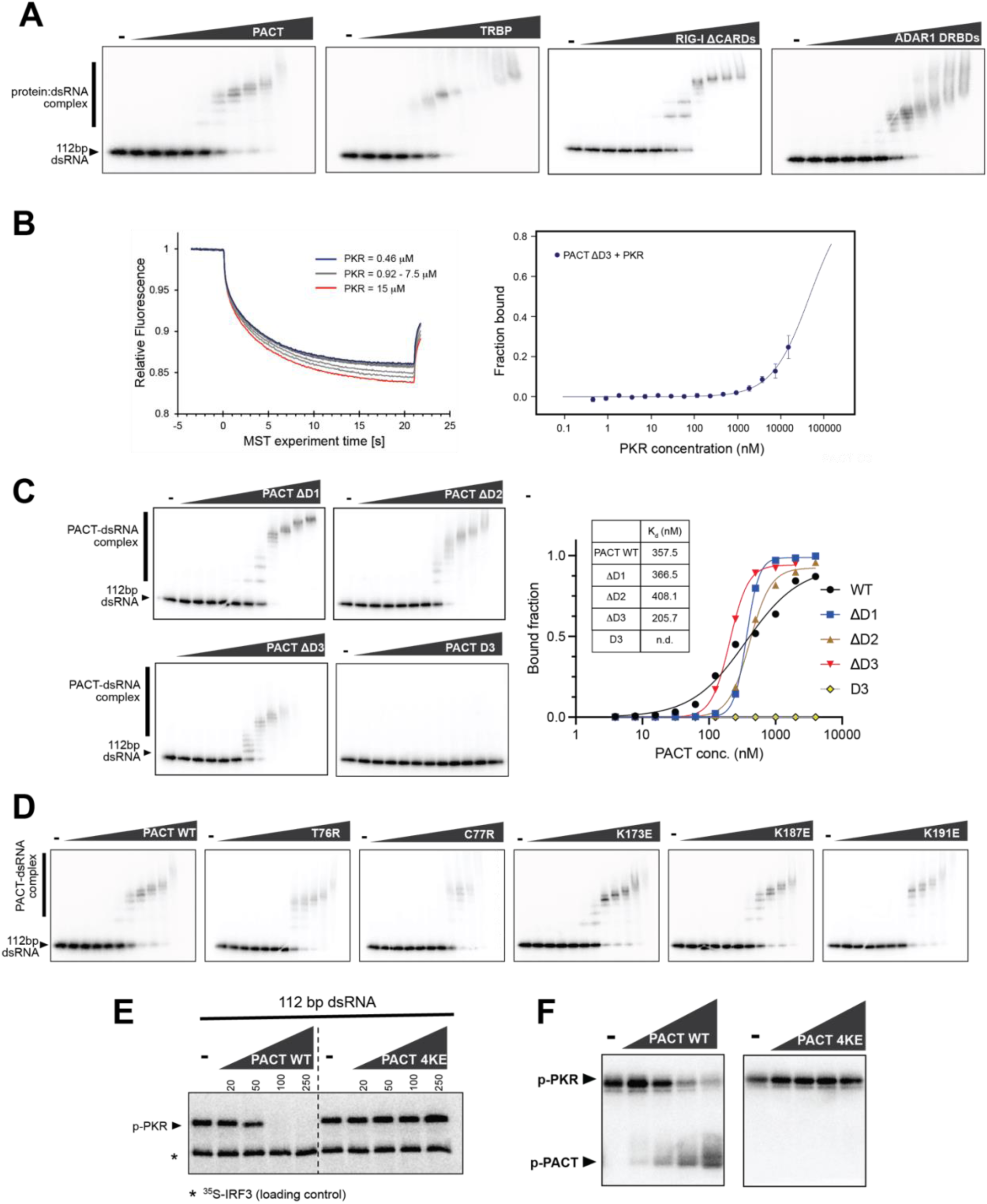
PACT forms a direct interaction with PKR to inhibit its kinase activity. (A) Native gel-shift assay monitoring 112 bp dsRNA binding to increasing concentrations (0, 3.9, 7.8, 15.6, 31.3, 62.5, 125, 250, 500, 1000, 2000, 4000 nM) of PACT, TRBP, RIG-I ΔCARDs, ADAR1 DRBDs. Binding curve analysis is shown in Figure 5B. (B) Microscale thermophoresis analysis showing binding of GST-tagged PKR (K296R) to FAM-labelled PACT ΔD3. MST trace (left) and dose-response curve (right) with PACT ΔD3 titrated against PKR (0.000458 to 15 μM), from 3 independent experiments. (C) Native gel-shift assay monitoring 112 bp dsRNA binding to increasing concentrations (0, 3.9, 7.8, 15.6, 31.3, 62.5, 125, 250, 500, 1000, 2000, 4000 nM) of PACT wild type, ΔD1, ΔD2, ΔD3, D3. Right: RNA binding curves. (D) Native gel-shift assay monitoring 112 bp dsRNA binding to increasing concentrations (0, 3.9, 7.8, 15.6, 31.3, 62.5, 125, 250, 500, 1000, 2000, 4000 nM) of PACT wild type, T76R, C77R, K173E, K187E, K191E. Binding curve analysis is shown in Figure 6B. (E) *In vitro* PKR kinase assay using 112 bp dsRNA (0.25 ng/μl) in the presence of 0, 20, 50, 100, 250 nM PACT WT and 4KE. (F) RNA-independent PKR kinase assay with 1.5 μM PKR in the presence of 0, 1.25, 2.5, 5, 10 μM PACT WT and 4KE. Based on our results in Figure 6, where mutations in β3 of DRBD2 impair PACT functions while equivalent mutations in β3 of DRBD1 do not, we speculate that K84 and K85 of DRBD1, together with β3 of DRBD2, are involved in PKR binding. Consistent with this hypothesis, 4KE does not undergo phosphorylation by PKR, unlike WT PACT, suggesting an impaired PKR-PACT interaction.

**Figure S7.**
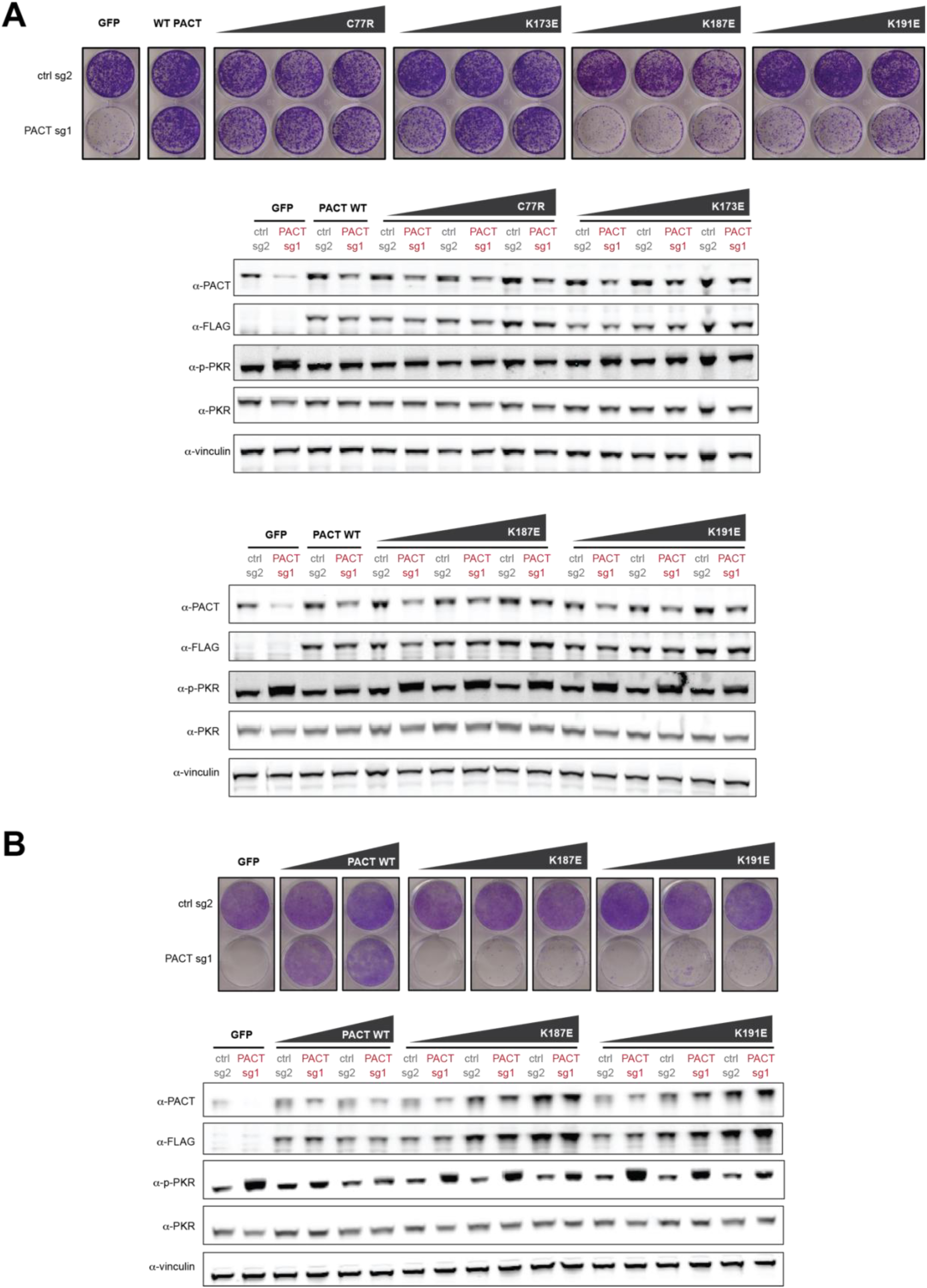
PACT DRBD2 interface for PKR interaction is critical for PKR suppression. (A) PACT complementation assay using crystal violet staining and Western blot analyses. NCI-H727 cells, either control gene-KO and PACT-KO, were complemented with increasing amounts of lentiviruses expressing GFP, Flag-tagged PACT wildtype, C77R, K173E, K187E or K191E. (B) PACT complementation assay using crystal violet staining and Western blot analyses. HCC1806 cells, either control gene-KO and PACT-KO, were complemented with increasing amounts of lentiviruses expressing GFP, Flag-tagged PACT wildtype, K187E or K191E.

**Supplementary Table S1.** NMR and refinement statistics.

| **NMR distance and dihedral constraints<sup>a</sup>** | **A (30-99)** | **B (30-99)** |
| --- | --- | --- |
| Distance constraints from NOE | 202 | 198 |
| Short-range intramolecular ($\vert i-j\vert \leq 4$) | 139 | 145 |
| Long-range intramolecular ($\vert i-j\vert \geq 5$) | 38 | 28 |
| Intermolecular | 25 |  |
| Total dihedral angle restraints<sup>b</sup> | 110 | 112 |
| $\phi$ (TALOS) | 55 | 56 |
| $\psi$ (TALOS) | 55 | 56 |
| Hydrogen bond restraints |  |  |
| Intramolecular | 18 | 18 |
| Intermolecular | 6 |  |
| **Structure statistics<sup>c</sup>** |  |  |
| Violations (mean ± s.d.) | | |
| Distance constraints (Å) | 0.062 ± 0.004 | |
| Dihedral angle constraints (°) | 0.564 ± 0.028 | |
| Deviations from idealized geometry |  |  |
| Bond lengths (Å) | 0.005 ± 0.000 | |
| Bond angles (°) | 0.626 ± 0.008 | |
| Impropers (°) | 0.466 ± 0.018 | |
| Average pairwise r.m.s. deviation (Å)<sup>d</sup> |  |  |
| Heavy | 1.599 |  |
| Backbone | 0.985 |  |

<sup>a</sup> The numbers of constraints are summed over two subunits.
<sup>b</sup> Backbone $\phi$ and $\psi$ restraints and their respective uncertainties were obtained from the “GOOD” dihedrals generated by the TALOS+ program based on the backbone chemical shift values.
<sup>c</sup> Statistics are calculated and averaged over an ensemble of the 15 lowest energy structures out of 75 calculated structures.
<sup>d</sup> The precision of the atomic coordinates is defined as the average r.m.s. difference between the 15 final structures and their mean coordinates except the flexible loop region (residue 52 to 59).

**Supplementary Table S2.**
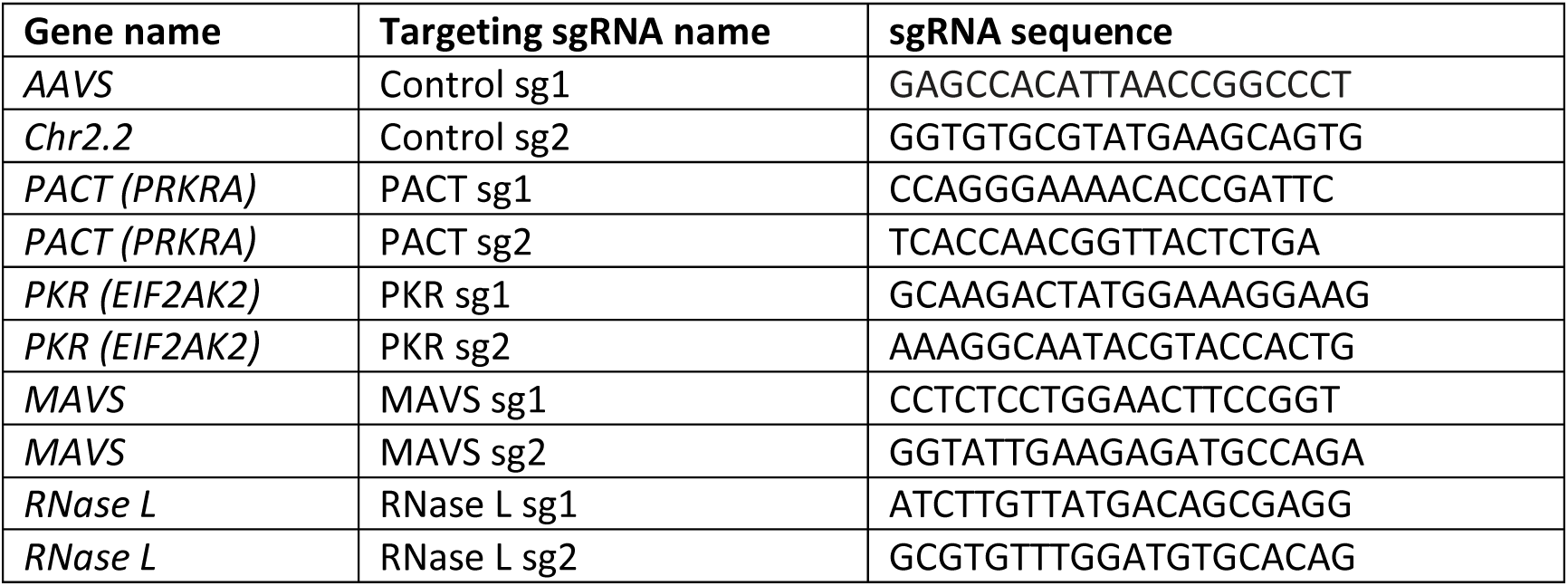
sgRNA sequences utilized for CRISPR-Cas9-mediated gene deletion. The targeted gene (column A), name of the sgRNA used in this study (column B), and sgRNA sequence (column C) are provided.

**Supplementary Table S3.**
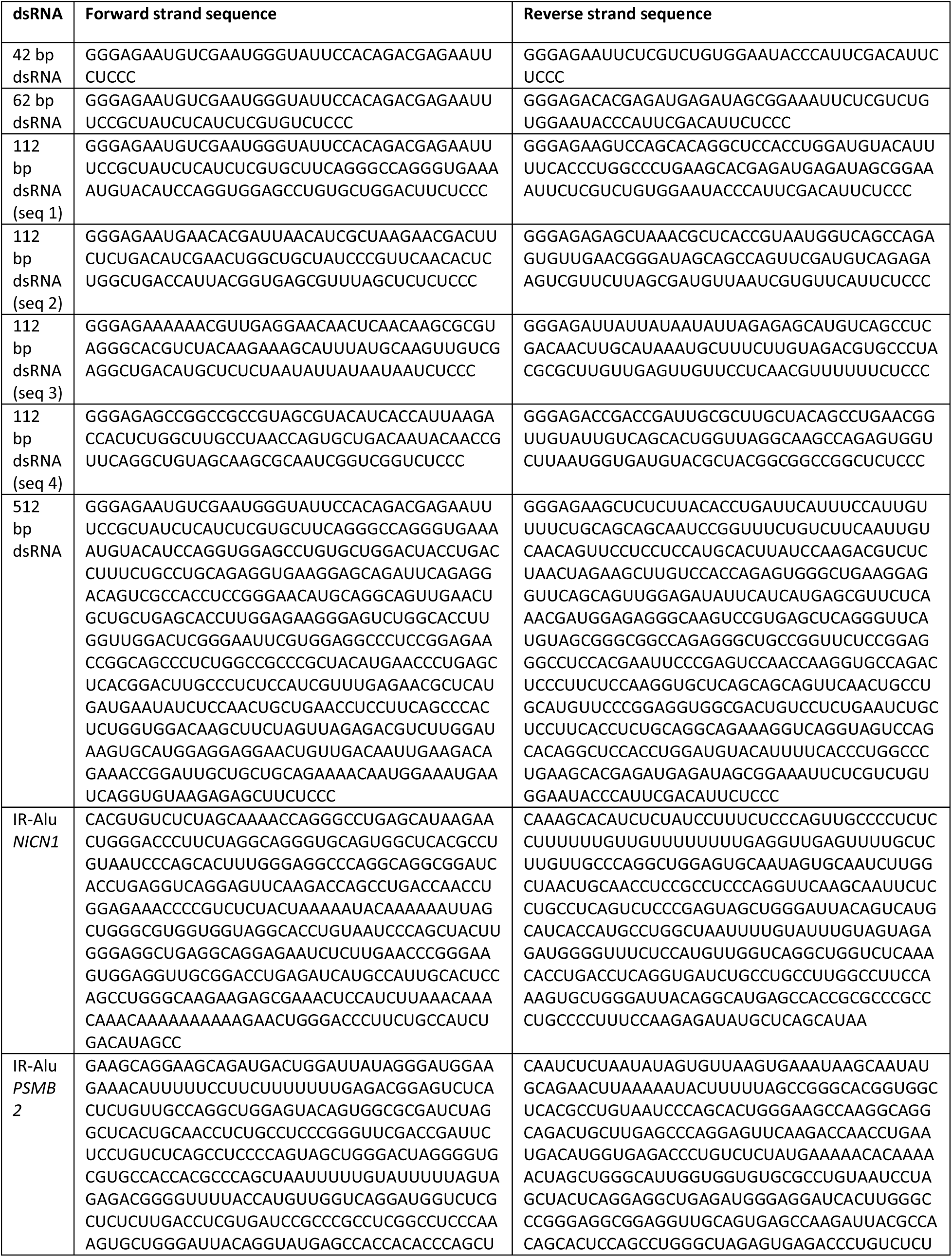
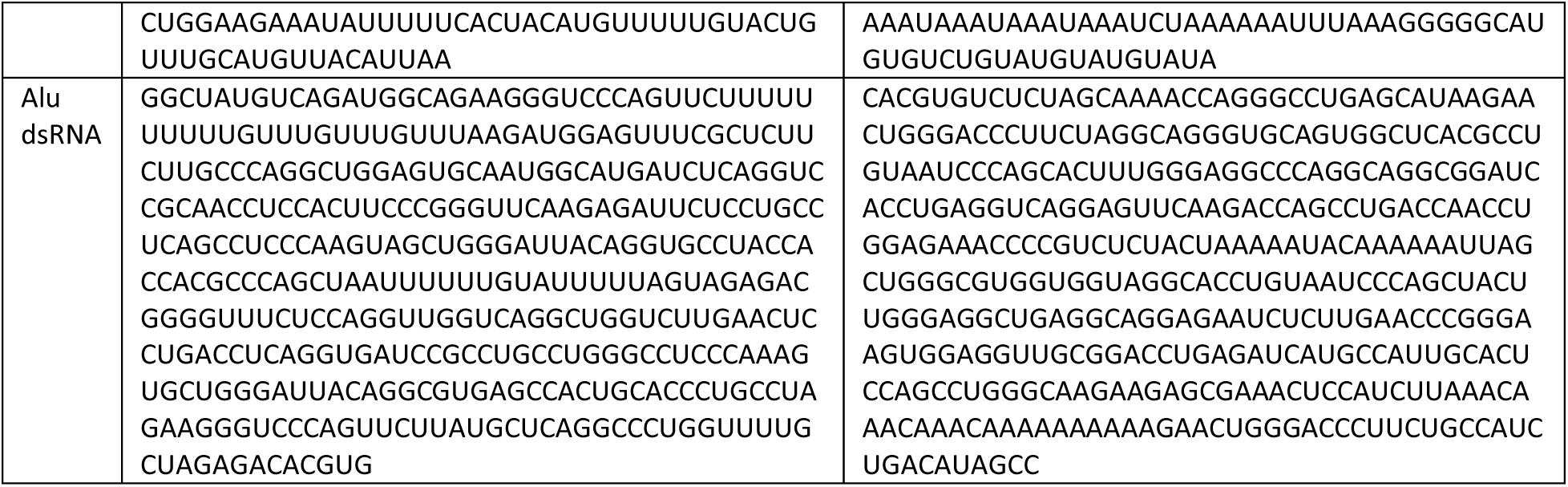
Sequences of dsRNAs used in the study.

## Notes

### Competing Interest Statement

The authors have declared no competing interest.

